# How grazing and other disturbance factors influence biodiversity in Cyprus, with a focus on breeding bird communities

**DOI:** 10.1101/860296

**Authors:** M. A. Hellicar, A. N. G. Kirschel

## Abstract

Grazing and browsing by sheep and goats has been an important anthropogenic influence on ecosystems in the Mediterranean region for centuries. This influence has changed significantly in recent decades, with a general shift from free-range grazing to the penning of animals. The intermediate disturbance hypothesis (IDH) proposes that perturbation - including anthropogenic disturbance - is the “norm” for ecosystems, and Mediterranean systems in particular, and that higher species diversity is found under conditions of continuous, low-level perturbation. We used Cyprus as a case study with the aim of assessing the impact of changes in grazing practice on biodiversity, while also taking account of other anthropogenic factors, such as fire. We aimed to test the IDH as it relates to grazing of scrub and open forest habitats in Cyprus, in the context of the general shift away from free-range grazing. Our hypothesis was that a greater diversity of breeding birds and plants would be found at sites subject to continuous low-level grazing perturbation over a long period of time, compared to sites where grazing has recently ceased, overgrazed sites and sites that have never been grazed.

We carried out surveys of breeding birds and vegetation at 48 study sites in scrub and open woodland across Cyprus. We estimated relative grazing pressure (past and present) and fire history at these sites, and looked for associations between these factors and birds, and perennial vegetation.

Our findings showed the importance of anthropogenic disturbance for biodiversity in scrub and open forest habitats in Cyprus. However, our results relating to the influence of fire and grazing on birds and vegetation suggest that it is not a regime of continuous low-level disturbance, but rather the absence of perturbation – or at least only very low-level perturbation – that benefits biodiversity in these habitats in Cyprus. This suggests the best approach for biodiversity management in scrub and open woodland habitats in Cyprus is to keep grazing to a minimum and avoid fires, though a ‘no grazing’ approach should also be avoided, because absence of grazers would likely increase fire risk.

## Introduction

Grazing by domesticated sheep *Ovis aries* and grazing and browsing by domesticated goats *Capra aegagrus* has played a key part in shaping Mediterranean landscapes and wildlife habitats (Blondel & Aronson 1999, Grove & Rackham 2001, Allen 2001, Blondel *et al*. 2010). Grazing and browsing are long-established practices that have undergone significant change in recent decades in many parts of the Mediterranean, with a shift to stall-feeding of sheep and goats (Grove & Rackham 2001, Blondel & Aronson 1999, Harris 2007). This change in an important anthropogenic disturbance factor has potentially far-reaching consequences for habitats and therefore for biodiversity and its conservation. Taking the eastern Mediterranean island of Cyprus as our example, our study aims to investigate the effects of grazing and browsing pressure in the past and present on biodiversity. Our focus is on the interaction of grazing with breeding season bird abundance and diversity, but also on plant diversity. We also look at how plant diversity affects breeding birds, because this is a possible mechanism for indirect influence of grazing on the bird community. We also investigate the interaction of grazing with other factors natural and man-made, such as fire, geology and rainfall.

Along with the clearing of land for agriculture, fire and grazing are considered the dominant anthropogenic forces creating and maintaining typical semi-natural Mediterranean scrub habitats (phrygana, garrigue and maquis), that also influence forest structure (Blondel & Aronson 1999, Grove & Rackham 2001, Allen 2001, Harris 2007, Blondel *et al*. 2010). We follow Tomaselli (1977) in defining phrygana as open stands of sclerophylous bushes up to 60 cm tall, garrigue as open stands of taller bushes (60 cm – 2 m), and maquis as more dense stands of bushes over 2 m tall. Grasslands are another important habitat influenced by grazing (Blondel & Aronson 1999, Allen 2001, Blondel *et al*. 2010), but we do not focus on this vegetation type in our study, as it is a very rare habitat of limited distribution in Cyprus (Tsintides *et al*. 2007). Habitats shaped by human activity can be important for biodiversity, notably low-intensity cultivation (Bignal & McCracken 1996, Pain & Pienkowski 1997, Tucker & Evans 1997, Tscharntke *et al*. 2005, Stoate *et al*. 2009, Oppermann *et al*. 2012, Emmerson *et al*. 2016, Navarro & Bao 2018, Traba & Morales 2019), and scrub habitats created and maintained by grazing, especially in the Mediterranean (Tucker & Evans 1997, Blondel & Aronson 1999, Grove & Rackham 2001, Harris 2007, Oppermann *et al*. 2012, Blondel *et al*. 2010, Doxa *et al*. 2010, 2012, Ieronymidou 2012).

Grazing and browsing have a recognised and significant effect on biodiversity in the Mediterranean area in general, with most previous work having focused on the direct effects on vegetation (Naveh & Whittaker 1979, Montalvo *et al*. 1993, Noy-Meir 1998, Papanastasis 1998, Peco *et al*. 1998, Papanastasis, Kyriakakis & Kazakis 2002, Shachak *et al*. 2008, Arga & Ne’eman 2009). Goat browsing has the greater impact on perennial vegetation and therefore on vegetation structure and composition, whereas sheep need grass and herb pasture to feed on and have little capacity to transform established woody vegetation (Blondel & Aronson 1999, Blondel *et al*. 2010). Hereafter, we use the term ‘grazing’ to refer to the combined effects of grazing and browsing.

Domesticated goats and sheep act as ‘allogenic ecosystem engineers’, modifying the habitats they graze by shifting the availability of resources for other organisms through their impact on vegetation and soil, and in particular through their impact on ‘landscape modulator’ plant species (Shachak *et al*. 2008, Arga & Ne’eman 2009). Grazing can generate habitat patchiness, helping to create a landscape mosaic of areas with differential availability of resources and thus varying distributions of organisms (Grove & Rackham 2001, Shachak *et al*. 2008, Arga & Ne’eman 2009, Blondel *et al*. 2010).

In the Mediterranean region, the history of human influence on the natural environment dates back more than 10,000 years (Allen 2001, Goren-Inbar *et al*. 2004, in Arga & Ne’eman 2009). With the start of agriculture about 10,000 years ago and its spread across the Mediterranean region between 10,000 and 5,700 years ago (Allen 2001), humans begin to have a big impact on natural habitats, predominantly through tree clearing and grazing (Blondel & Aronson 1999, Grove & Rackham 2001, Allen 2001, Blondel *et al*. 2010). Domestication of sheep and goats took place between 9,000 and 8,000 years ago (Allen 2001). In recent decades, grazing practices have undergone much change in the Mediterranean area, with a widespread move towards penning of animals and a consequent reduction in free-range grazing (Blondel & Aronson 1999, Grove & Rackham 2001, Blondel *et al*. 2010). This change has potentially far-reaching consequences for biodiversity, because it lessens or even removes the influence of ecosystem engineer species (Shachak *et al*. 2008, Arga & Ne’eman 2009).

Grazing, especially when combined with fire, can substantially reduce tree cover by arresting natural vegetation succession processes (Grove & Rackham 2001, Allen 2001, Papanastasis, Kyriakakis & Kazakis 2002). Reduction of perennial vegetation cover will tend to increase soil erosion rates, as exposure to precipitation and wind is increased, with vegetation cover of 65–70 % needed to prevent soil loss due to erosion (Duran Zuazo & Rodriguez Pleguezuelo 2008). While grazing can be incompatible with forestry policies aiming at maximising forest cover (Thirgood 1987), more recent work challenges the traditional conservationist view that grazing degrades ecosystems (Grove & Rackham 2001, Harris 2007, Blondel *et al*. 2010). Indeed, it has been shown that Mediterranean ecosystems are often resilient when grazed – even intensively - and that limitation or cessation of grazing can lead to biodiversity loss (Grove & Rackham 2001, Allen 2001, Papanastasis, Kyriakakis & Kazakis 2002, Blondel *et al*. 2010). But this resilience does not equate to tolerance to the overgrazing that affects many Mediterranean habitats (Giourga *et al*. 1998, Sales-Baptista *et al*. 2016)

Grove & Rackham (2001) and Allen (2001) challenge the traditional view that the whole Mediterranean basin was once covered in climax forest vegetation and is now in a ‘degraded’ state due to the impact of forest clearance, fire and grazing. Grove & Rackham (2001) state that the reality is far more complex: Mediterranean vegetation has come to be dominated by grazing and fire-resistant species in a community that has evolved resistance to the thousands-of-years-long influence of human cutting, burning and especially grazing activities and forest is not always the natural climax community. In the Mediterranean basin, grazed maquis, garrigue and phrygana communities often support higher species diversity than forest (Grove & Rackham 2001, Papanastasis 2004). Grazing, and even tree felling and controlled burning, generate a regional mosaic (Grove & Rackham 2001, Blondel *et al*. 2010) rich in landscape-scale biodiversity or γ-diversity (Whittaker, 1960). Allen (2001) states that while grazing will keep vegetation low in scrub communities, traditional, moderate and rotational grazing encourages a productive and diverse herbaceous plant community. Lifting of grazing pressure can allow grasses and thistles to dominate such systems, reducing diversity, while overgrazing will lead to unpalatable species dominating, which can increase fire risk as fuel builds up in the system and many of these unpalatable species are fire-adapted. Use of fire as a traditional management tool in grazed habitats can restrict unpalatable woody vegetation and encourage more palatable plants, and help in the maintenance of an equilibrium between livestock numbers and scrubby habitats. But cessation of grazing can also lead to sites becoming more fire-prone and more prone to intense fires, because of the accumulation of fuel, particularly in the form of dead vegetation (Blumler 1996, Grove & Rackham 2001, Allen 2001, Papanastasis, Kyriakakis & Kazakis 2002).

The intermediate disturbance hypothesis (IDH) proposes that higher species diversity (in particular for local scale or α-diversity) is found in ecosystems where disturbance is neither too rare nor too frequent (Wilkinson 1999). The IDH proposes that systems that have evolved with a regime of continuous perturbation require disturbance through fire, grazing, wood cutting and other - largely but not exclusively anthropogenic - actions to maintain their diversity (Wilkinson 1999, Blondel & Aronson 1999, Allen 2001, Blondel *et al*. 2010).

Plants can respond to grazing through the evolution of tolerance or avoidance mechanisms. Tolerance mechanisms include compensatory growth and enhanced fecundity, while avoidance mechanisms can involve accumulation of secondary metabolites, morphological and phenological adjustments (Dobarro, Valladares & Peco, 2010). Mediterranean plants have also evolved tolerance mechanisms for coping with periodic fires, such as the capacity to put out fresh shoots from the roots (Blondel & Aronson 1999, Allen 2001). The frequency and severity of fires has been shown to be important in its impact on vegetation, and in relation to recovery after fire in particular (Fernández-García *et al*. 2019).

A number of studies have looked at the interaction between grazing and biodiversity in the Mediterranean, mostly in Greece, Israel or Spain (Naveh & Whittaker 1979, Montalvo *et al*. 1993, Noy-Meir 1998, Papanastasis 1998, Peco *et al*. 1998, Papanastasis, Kyriakakis & Kazakis 2002, Shachak *et al*. 2008, Arga & Ne’eman 2009). The focus of these studies has been on the grazing impact on vegetation. Papanastasis (2004), in his review on past and present environmental management practices on Crete, found that traditional, low-intensity management by farmers, shepherds and foresters created a diverse landscape mosaic rich in biodiversity, with intensification of these activities, particularly after World War II, having had generally negative results for biodiversity. Intermediate disturbance has also been found to increase plant biodiversity in maquis vegetation in Israel (Carmel & Kadmon 1999, Shachak *et al*. 2008 and Arga & Ne’eman 2009).

Heavily grazed phrygana sites in Crete were found to have higher species richness and diversity than protected - grazing free – sites. This pattern did not hold where there was burning combined with heavy grazing, with such sites having lower species richness and lower diversity than sites that were only heavily grazed (Papanastasis, Kyriakakis & Kazakis, 2002). Burning reduced the cover of chamaephytes, which removed from the system the protection from grazing these afford for therophytes and geophytes (Papanastasis, Kyriakakis & Kazakis 2002). Fire was found to have a rejuvenating effect on the phrygana community, encouraging herbaceous growth and attracting overgrazing (Papanastasis, Kyriakakis & Kazakis, 2002). There can also be an impoverishment of the seed bank in areas where fire is combined with heavy grazing.

Work on Mediterranean grasslands in Israel (Naveh & Whittaker 1979, Noy-Meir 1998), also supports the importance of IDH, with plant diversity found to be greater under moderate grazing than when grazing pressure was removed. Only extreme overgrazing (such as around watering holes) was found to actually reduce biodiversity (Noy-Meir 1998). For Mediterranean grasslands in Spain, Montalvo *et al*. (1993) found that species diversity had declined four years after cessation of grazing, while Peco *et al*. (1998) found a similar pattern for Dehesa grassland habitats. But patterns of enhancement of biodiversity in grasslands as a result of grazing have not been found in Greece, though it should be noted that the relevant case studies involve very high grazing pressure (Papanastasis, Kyriakakis & Kazakis 2002).

For grasslands in the Mediterranean, Noy-Meir (1998) found that there is an optimal, moderate, but also varying grazing pressure that maintains more species than either heavy grazing or no grazing. Where uniform grazing pressure is implemented across a landscape, there is a slow loss of species as a result of the gradual retreat and eventual disappearance of those species that are adapted to either heavy grazing or no grazing (Noy-Meir 1998). This makes grazing a difficult tool to use for habitat management for conservation, for while this optimal grazing regime often exists under traditional, unmanaged systems, it is hard to replicate in a more managed system. Papanastasis, Kyriakakis & Kazakis (2002) note that a number of studies have suggested that it is not just grazing pressure (stocking levels) but also grazing regime (the timing of grazing) that has an important influence on how grazing interacts with biodiversity.

Turning to Cyprus, archaeological evidence suggests cultivation and herding reached the island between 9,000 and 7,800 years ago (Allen 2001). In keeping with the pattern for the rest of the Mediterranean, grazing practices have changed in Cyprus in recent decades, with a move away from extensive goat and sheep grazing towards stall feeding of penned animals (Christodoulou 1959, Economides 1997, Harris 2007, FAO 2010, MARDE 2017). Historically, free-ranging flocks of mostly goats were grazed on semi-natural scrubland and open woodland in Cyprus, but the introduction of the 1935 Goat Law regulated goat distribution, causing a decline in numbers by 33% between 1930 and 1975 (Economides 1997) and this trend continued in subsequent decades. Continuous data is not available, but there was a 24% drop in the number of sheep and goat farms between 2010 and 2016. Despite this, the number of sheep and goats remained high, at around 400,000 over the same period, with sheep outnumbering goats by around two-to-one (MARDE 2017). Data for the numbers of free-range goats and sheep is not available after 1960, but there has been a clear shift in husbandry towards keeping animals penned and stall-fed, with free-range grazing more limited (Christodoulou 1959, Economides 1997, MARDE 2017).

A pattern of general decline in free-range grazing, while in some parts of Cyprus free-range grazing practices continue, sometimes intensively (Eliades *et al*. 2016), makes the island a good case-study for examining the effects of shifting grazing patterns on biodiversity. Cyprus is rich in both plant and avian diversity, with 108 endemic plant species (Tsintides 1995) and three endemic bird species (Flint & Stewart 1992, Flint *et al*. 2015) while the island also has significant breeding populations for a number of bird species considered to be of conservation priority on a European scale (Flint & Stewart 1992, BirdLife International 2004). The importance for bird diversity of a ‘patchy’ landscape with varying land uses has been shown for Cyprus (Ieronymidou, 2012) as it has been for Europe in general and the Mediterranean in particular (Bignal & McCracken 1996, Blondel & Aronson 1999, Benton *et al*. 2003, Blondel *et al*. 2010, Oppermann *et al*. 2012, Emmerson *et al*. 2016, Navarro & Bao 2018).

Regarding grazing and its impact on birds, Ieronymidou *et al*. (2012) suggest that a decline in grazing intensity will have caused a structural and successional shift from compact, tightly grazed dwarf-shrub phrygana to taller, open-structured garrigue and maquis in Cyprus, as has occurred in Crete (Papanastasis & Kazaklis 1998). This change is potentially significant for a number of bird species of conservation priority in Cyprus, such as Linnet *Linaria cannabina*, European Roller *Coracias garrulus*, Cretzschmar’s Bunting *Emberiza caesia*, Cyprus Wheatear *Oenanthe cypriaca*, and especially Cyprus warbler *Sylvia melanothorax*.

For the Mediterranean, intermediate disturbance is the regime many species and communities have evolved with and the intensification of cultivation and grazing practices (abandonment of free-range grazing) seen in recent years could have a detrimental effect on biodiversity (Attenborough 1987, Blondel & Aronson 1999, Allen 2001, Blondel *et al*. 2010). Allen (2001) states that disturbance is the “norm” for ecosystems, and Mediterranean systems in particular, and may be necessary for the continued existence of many communities and some species. Our aim in this study is to use Cyprus as a test case to investigate the validity of the IDH with regards to grazing primarily, but while also taking into account other key disturbance factors, notably fire. We hypothesise that for Cyprus, a greater diversity of breeding birds and a greater diversity of plants will be found where habitats have been subject to continuous low-level grazing perturbation over a long period of time, compared to sites where grazing has recently ceased, overgrazed sites and sites that have never been grazed.

## Methods

### Sampling Site Selection

Our sampling strategy aimed at capturing a wide variety of natural and semi-natural scrub and pine forest habitats that were subject to a range of current and past grazing pressures. As no systematic data on grazing pressure was available, we opted for a simple, two-step site selection approach that aimed to maximise variation in coverage of relevant habitats:

i. We used Coordination of information on the environment (CORINE) programme land-cover data (https://www.eea.europa.eu/publications/COR0-landcover) to define six study areas dominated by phrygana, garrigue, maquis and pine forest habitat. Using the 2006 CORINE Land-cover (CLC) map for Cyprus (the most up-to-date available at the time), we selected the following CLC codes: 312 ‘Coniferous forest’, 321 ‘Natural grasslands’, 323 ‘Sclerophyllous vegetation’, 324 ‘Transitional woodland shrub’ and 333 ‘Sparsely vegetated areas’ to capture the phrygana to forest habitat spectrum. We used the 2000 and 2006 Cyprus CLC maps to exclude all recently burnt areas (CLC code 334). Each study area had a uniform underlying geology. The study areas were (geology in parentheses): Akamas Peninsula (Mamonia complex), Xeros/Diarizos valleys (Marls, Chalk & Clays), West Limassol district (Chalk), West Larnaca district (Chalk), Southeast Nicosia district (igneous including pillow lavas) and Karpasia Peninsula (Sandstone-Gypsum).
ii. We explored each study area to identify the dominant scrub/forest vegetation communities and to select 48 sampling sites to capture these in a balanced design. A key selection criterion was accessibility and availability of tracks or paths for survey work. We ensured survey sites were widely spread across each study area and were a minimum of 1 km apart. Site altitude ranged from close to sea level to just over 700 m, with two sites above 1000 m.

For logistical reasons, the final site list included twelve sites in Larnaca and Nicosia and six sites in each of the more remote study areas: Akamas, Xeros/Diarizos, Limassol and Karpasia (Figure 2).

**Figure 1.**
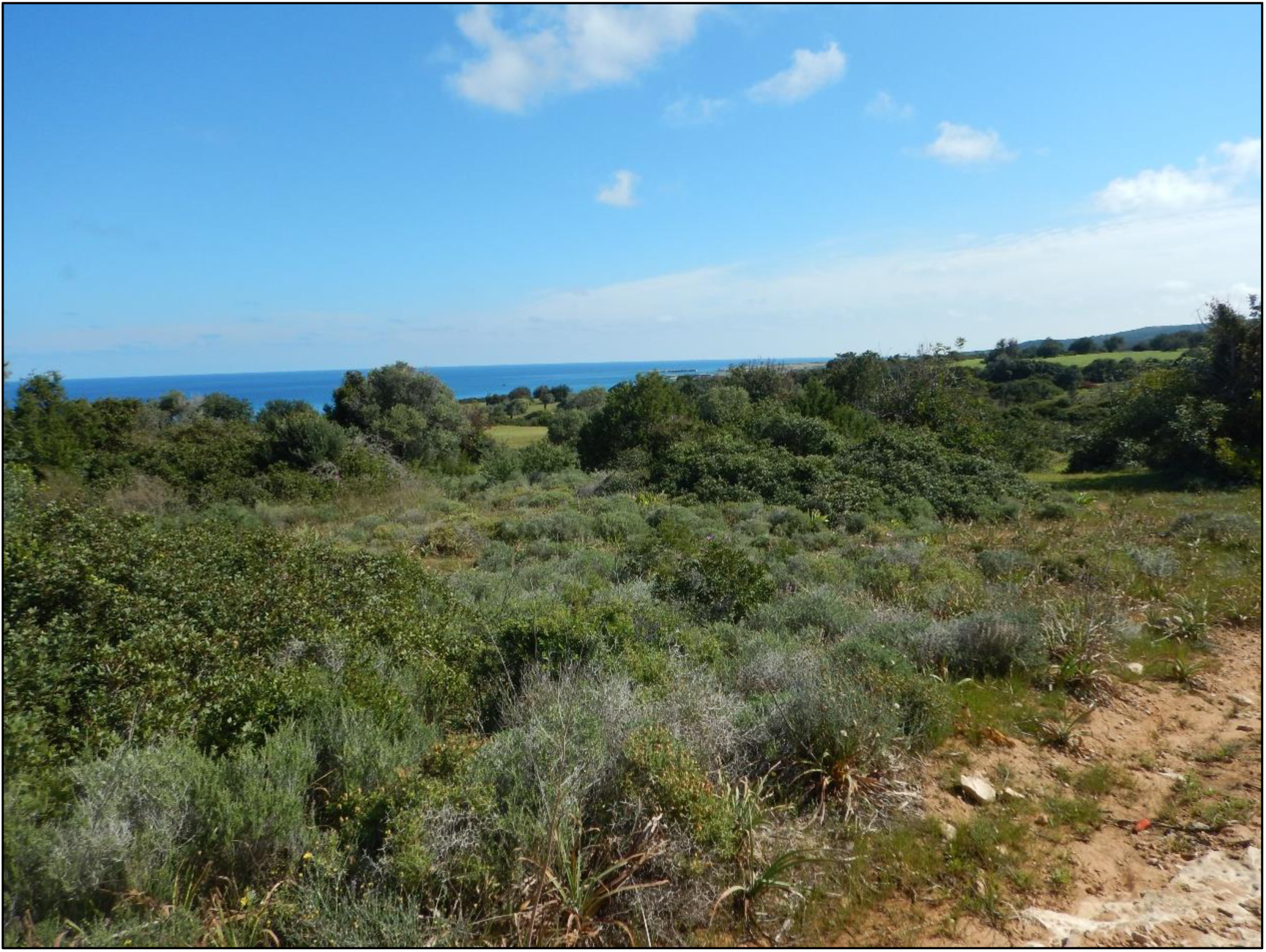
Grazed scrub in the Karpasia area (*photo by Athina Papatheodoulou*)

**Figure 2.**
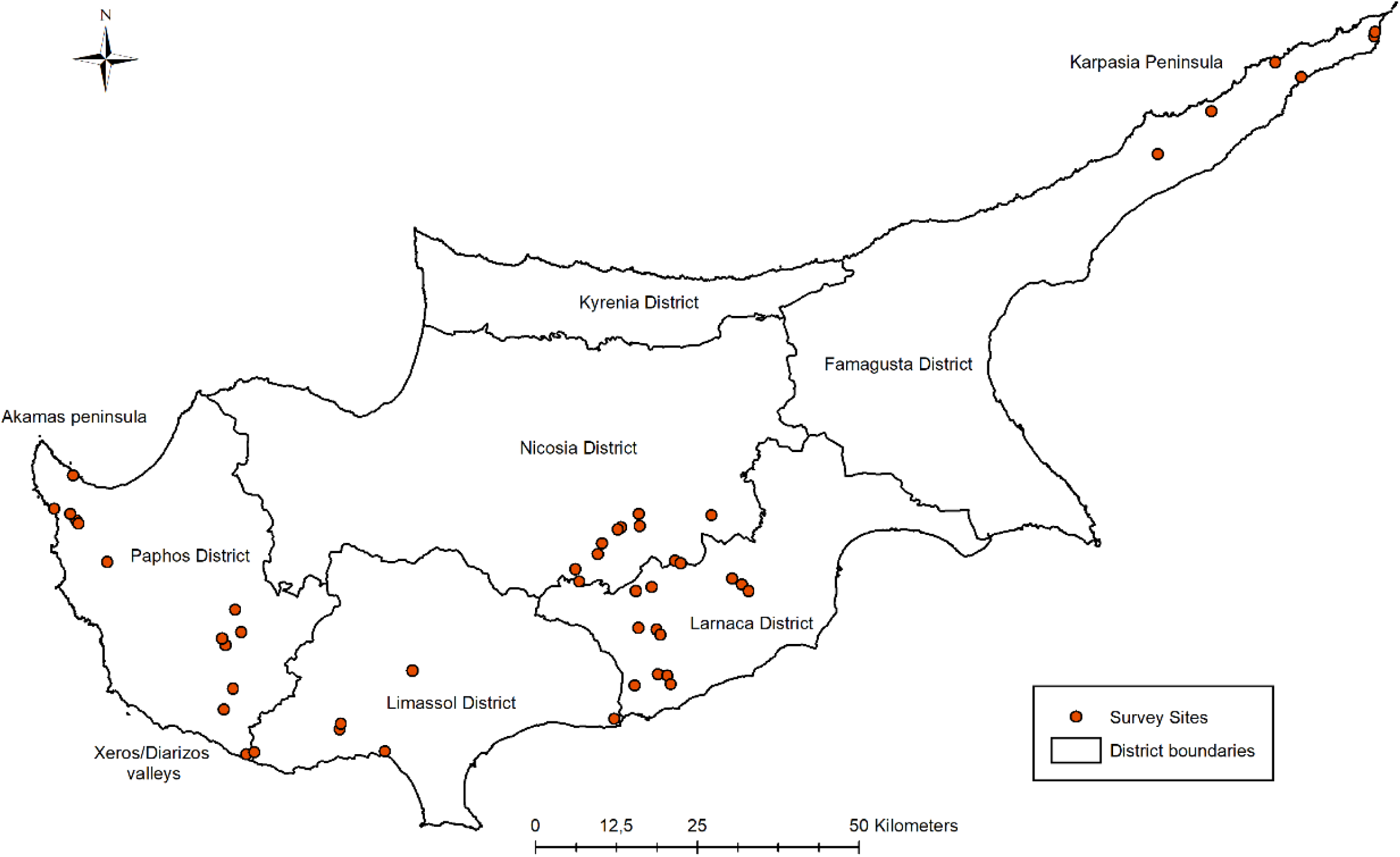
Location of the 48 survey sites across Cyprus

The flow diagram below shows how study site selection was arrived at:

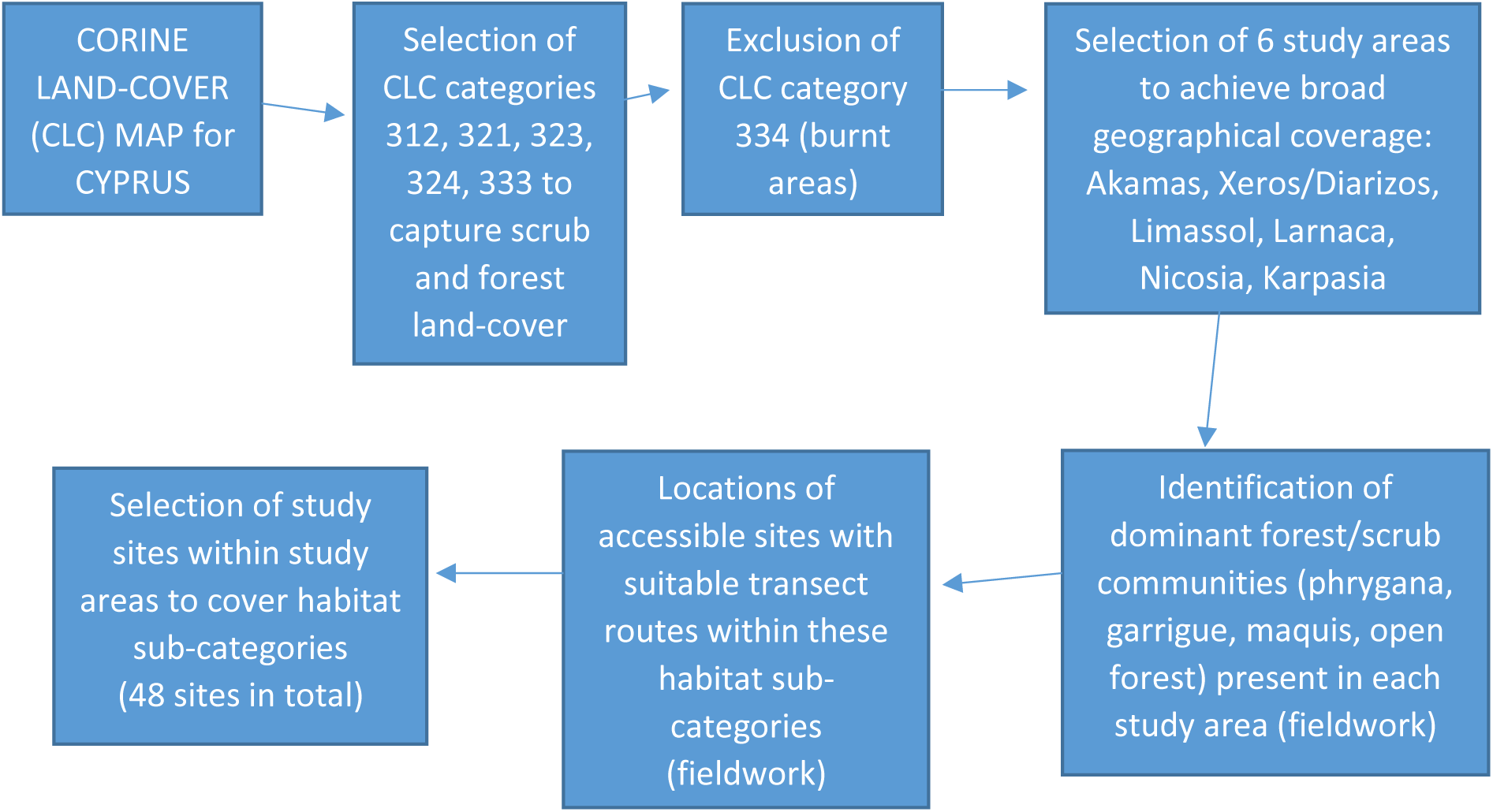

### Bird and vegetation surveys

#### Bird Surveys

The line transect method (Bibby *et al*. 2000) was used for bird surveys carried out (one transect per survey site) during the 2012, 2014 and 2016 breeding seasons. We covered 48 survey sites in total, but only 33 of these sites were covered in 2012. Survey sites were visited twice each year, once early and once late in the breeding season. Early season visits were from mid-March to end of April, to cover the peak period of song and display (and therefore peak detectability) of species such as *Sylvia* warblers, while late season visits were in May and June, a better period for detecting late-arriving migrant breeders such as European Roller and Black-headed Bunting *Emberiza melanocephala*. Early and late visits to any given site always took place at least two weeks apart. Transects mostly followed dirt tracks though some were off-track, at least in part. They averaged 825 m in length, while generally varying from 500 m to 1200 m, with one 3350 m long. The survey protocol was to walk transect routes at a slow pace while recording all birds seen or heard in four distance intervals: 0-10 m, 10-25 m, 25-50 m and 50-100 m from the transect line (distances were checked using a Bushnell Medalist laser range-finder). Surveys were completed in the four hours after sunrise and were done on days with wind-free and rain-free conditions. The recorder never walked directly towards the sun. Especially in more densely vegetated maquis and garrigue sites, most bird recordings were first made based on vocalisations, usually followed by visual confirmation.

#### Vegetation Surveys

Vegetation surveys were carried out at the 48 study sites using the same transect routes followed for bird transects. The vegetation surveys were completed in the winter and early spring (January to March) of 2017, with the aim of estimating cover and species diversity for perennial plants within the area surveyed for birds. The vegetation surveys also aimed to record field evidence relating to grazing pressure, including the presence of tracks made by goats and sheep, goat and sheep droppings and the presence of gazing-tolerant plant species associated with grazed habitats.

We used the frame quadrat method (Krebs 1999, Sutherland, 2000) to record perennial species and estimate cover for each one. A 2 x 2 m quadrat was laid down at twelve equally spaced points along the transect line. We alternated between laying the quadrats on the right and left of the transect line, and also in placing the quadrat at a distance of 0 m, 5 m, 15 m and 25 m from the line. At each quadrat sampling point, we identified all perennial species within the quadrat and estimated cover for each species, by eye, in three height bands: ground level - 60 cm, 60cm – 2m and > 2 m. The ground cover of herbaceous plants and grasses was also estimated, as was the cover of bare ground and stones/rock. Though these visual estimates of cover had an element of subjectivity, we assumed this would not vary significantly between sites as all estimates were made by the same observer.

The number of goat and/or sheep tracks crossing each 2 x 2 m quadrat was recorded and we counted droppings visible within each quadrat. We also recorded a measure of bush ‘compactness’, by placing a 25 x 25 cm black-and-yellow ‘checkerboard’ card at ground level behind the bush closest to the bottom-right-hand-corner of each quadrat and recording the proportion of the 25 squares visible from the other side of the bush (looking ‘through’ the plant). We recorded the following additional evidence relating to grazing at sites: the presence of obvious ‘browse lines’ on trees and larger bushes, the presence of grazing animals and of active goat pens/sheep-folds in the vicinity.

We also used the Point-Centred Quarter Method (PCQM, Cottam & Curtis 1956, Bullock 2000, Mitchell 2015) to estimate the density of trees and bushes over 2 m tall in a wider area including the transect and up to 500 m around this. At 15 randomly chosen points a minimum of 50 m apart along the transect line, we used a Bushnell Medalist laser range-finder and an electronic (smartphone) compass to record the distance to the nearest tree (or bush > 2 m tall) within each quadrant (NE, SE, SW and NW). The species of each tree/bush was also recorded as part of the PCQM survey, while no individual tree/bush was included more than once.

### Defining past and present grazing pressure & other explanatory factors

#### Estimating historical and current grazing pressure

Quantified, geographically referenced data on **past grazing pressure** was not available for Cyprus. We therefore used the best available indicator for this: the categorization of grazing regime under the 1935 Goat Law. At the time, and having already banned flocks from the State Forest areas established from 1879 onwards (Thirgood 1987), the British colonial administration introduced the Goat Law with the aim of restricting free-range goat grazing in areas beyond the state forests. Under the provisions of the 1935 law, village communities could decide (by a two-thirds majority vote) to become a ‘prescribed village’ where free range goat grazing was banned (Thirgood, 1987, Harris 2007). Despite a number of amendments, the basic provisions of the Goat Law were still in effect at the time of this study.

We used a 1982 Forestry Department Forest Map, showing prescribed village areas under the Goat Law and State Forests, as a basis for generating a GIS map using ArcGIS 10.2 (ESRI 2009) – see Figure 3. We assumed polygons corresponding to prescribed village areas and to state forests represented areas where there was likely to have been lower historical grazing pressure since the 1930s. While this regulatory regime does allow for exemptions under license and has not always been strictly adhered to in all areas - Akamas, for example, is a state forest area where free-range goat grazing persists (Eliades *et al*. 2016) - it represents the best available baseline for historical grazing patterns. The law’s restrictions do not apply to sheep and ‘prescribed’ village areas are therefore not absolute ‘no grazing’ areas.

**Figure 3.**
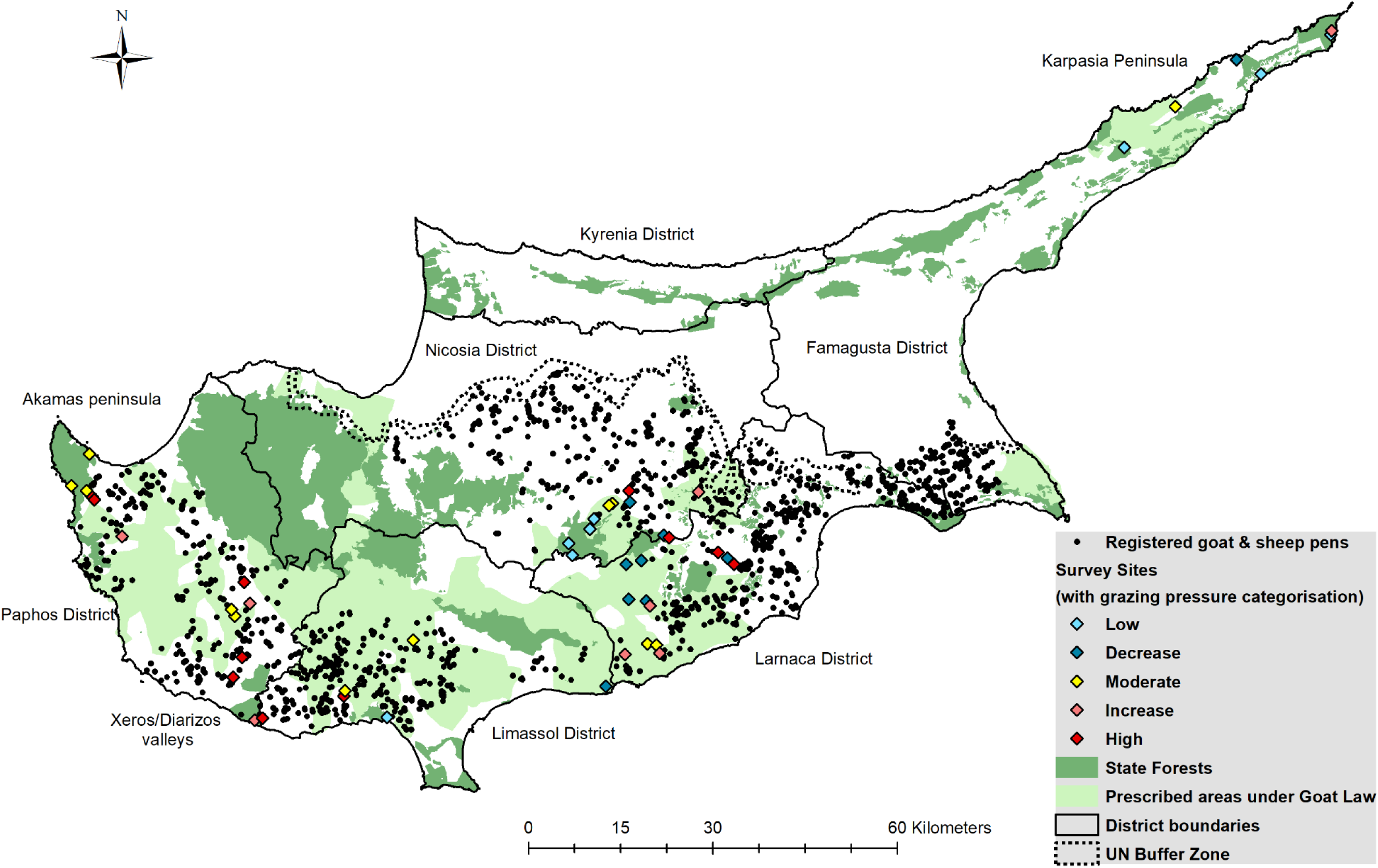
Grazing pressure overview map of Cyprus

Using this mapping exercise, we arrived at a broad, three-way categorization of sites for historical grazing pressure:

1. ‘LOW’: State Forest areas
2. ‘MODERATE’: areas prescribed under the Goat Law
3. ‘HIGH’: all other areas.

To estimate current grazing pressure, we used Cyprus Veterinary Service data on licensed goat pens and sheepfolds for 2012. This data includes the locations and numbers of animals at animal farms across Cyprus, but does not differentiate between goats and sheep. Based on Veterinary Service advice and information gained from informal interviews with shepherds in the field, we excluded farms with fewer than 25 animals or more than 750 animals, as such very small or very large pens tend to keep animals in all year round. This left 1,806 pens with 25 - 750 animals that we assumed were at least partly free-ranging. The location of these pens is indicated in figure 3. Eliades *et al*. (2016) note that the number of licensed animals shown for pens in the Veterinary Service data does not always match the reality on the ground, and we also found this to be the case for a number of animal pens we were able to estimate herd size for during survey work.

In the field, we recorded any pens housing free-ranging grazing animals within the actual area covered by the bird surveys. The animals in these pens we assumed to be ‘high impact’ local herds. Using GIS analysis, we then delineated 1 km and 3 km radius buffers around the center of each study site and summed the number of licensed grazing animals found in farms within these buffers. We estimated current grazing pressure (CGP) for sites as follows:

***CGP*** *= (number of animals penned within site x 2) + (number of animals penned within 1km buffer) + (number of animals penned within 3 km buffer / 4)*

This estimation approach was arrived at after discussions with shepherds and Veterinary Service officials; it is based on the norms regarding distances free-ranging livestock move from pens.

For the area north of the UN buffer zone in Cyprus, no data on numbers of licensed animals was available, so a different approach to gathering this data had to be adopted for the Karpasia study area. We used a combination of work in the field and examination of Google Earth satellite maps and 1:50000 Ordinance Survey (OS) maps, to locate goat pens and sheepfolds within 3 km of survey sites and estimate the number of animals present in these. We added to this livestock total the number of feral donkeys *Equus africanus asinus* recorded during surveys, to account for additional grazing pressure from these large free-ranging grazers. This approach allowed us to arrive at a data set on goat pens and livestock numbers for Karpasia that was comparable to the data set provided by the Veterinary Services for Cyprus South of the UN buffer zone.

Based on CGP estimates, we arrived at a three-way categorization for current grazing pressure:

‘LOW’: sites with CGP score < 50,

‘MODERATE’: sites with 50 - 400 CGP score, and

‘HIGH’: sites with > 400 CGP score.

We then checked these categorizations against indicators of grazing pressure recorded on site. These were: tracks made by grazing animals; grazing animal droppings; bush ‘compactness’ (as an indication of browsing) and prevalence of indicator plant species associated with high grazing intensity: *Drimia aphylla*, *Asphodelus ramosus*, *Sarcopoterium spinosum*, *Rhamnus lycioides graeca, Thymus capitatus, Lithodora hispidula versicolor*, and *Genista fasselata fasselata* (based on Meikle 1977, 1985). In a minority of cases where this field data was inconsistent with the CGP categorization, we revised this categorization upwards or downwards accordingly.

Finally, we combined the categorizations for past and present grazing pressure to arrive at five overall grazing pressure categories, as follows:

1. ‘LOW’ grazing sites: consistently low (or absence of) grazing.
2. Grazing ‘DECREASED’ or ‘RELEASE’ sites: reduction of grazing pressure in recent decades.
3. ‘MODERATE’ grazing sites: consistently grazed over time, but not heavily
4. ‘INCREASED’ grazing sites: raised grazing pressure in recent decades
5. Overgrazed or ‘HIGH’ grazing sites: areas consistently grazed, heavily

The matrix below shows how categorizations for past and present grazing pressure were combined to arrive at the overall grazing pressure categories:

|  |  | Historical Grazing pressure |  |  |  |
| --- | --- | --- | --- | --- | --- |
|  |  | LOW<br>(State Forest area) | MODERATE<br>(Prescribed village area) | HIGH<br>(other areas) |  |
| Current Grazing Pressure | LOW<br>(CGP <50) | LOW | DECREASED /<br>RELEASE | DECREASED /<br>RELEASE | Overall Grazing Pressure Category |
|  | MODERATE<br>(CGP 50-400) | INCREASED | MODERATE | DECREASE /<br>RELEASE |  |
|  | HIGH<br>(CGP >400) | INCREASED | INCREASED | HIGH |  |

Table 1 (in Appendix) shows the grazing data and grazing pressure categorization for all study sites.

While our focus was on grazing pressure, we also accounted for potentially confounding explanatory factors either by controlling for them experimentally or by quantifying them for inclusion in our statistical models.

#### Categorization for site fire history

We excluded from site selection all areas burnt in the 12 year period before our study began (2000-2011) by selecting against areas categorized as burnt (CLC code 334) on the 2002 and 2006 CORINE Land-cover (CLC) maps for Cyprus. There were no wildfires in the study site areas during the study period, 2012 - 2017. We double-checked this by examining Cyprus Department of Forests records for wildfires for the period 2010-2018 (downloaded from the National Open data portal https://www.data.gov.cy/node/47?language=en on 14.1.2019).

No systematic georeferenced data on wildfires was available for the period before 2000. Therefore, to determine if any survey sites (including the area within a 3km radius of sites) had burnt in the period between 1960 and 1999, we used the following sources:

i. Visual examination of Cyprus Lands and Survey Department (LSD) satellite images (http://eservices.dls.moi.gov.cy/#/national/geoportalmapviewer accessed on 15.1.2019) for 1963 and 1993 to identify evidence of bare or blackened areas that would point to fires in the periods around 20-30 years and 50-55 years before survey start. This approach was backed-up by examination of Google Earth imagery for Cyprus, which provided evidence dating back to 1984, though not for all study areas, with burnt areas visible as bare or blackened landscapes.
ii. Evidence of past burns in the field: presence of any burnt tree stumps or blackened remains of burnt bushes and/or presence of even-age stands of bushes such as *Cistus creticus, Cistus salviifolius*, *Genista fasselata fasselata*, *Calycotome villosa*, and *Sarcopoterium spinosum*. Such fire-tolerant shrubs will tend to experience a synchronous flush of growth after a fire event (Grove & Rackham 2010, Allen 2001, Blondel & Aronson 1999).

Based on the above, we coded sites as follows for fire history:

0. “Unburnt” sites – No evidence of fire for the past 50 years at least
1. Recovered sites – Evidence of older fire(s), approximately 20-55 years before survey start.
2. Recovering sites – Evidence of fire in the recent past, 12-20 years before survey start.
3. Burnt sites – Evidence of both older and more recent burns. Note there were no survey sites falling into this last category.

#### Definition of other potential explanatory variables

We defined other variables that could influence biodiversity at study sites:

i. Elevation: the average height above sea level for the transect route.
ii. Rainfall: the mid-point of the average annual range in rainfall for the period 1990 - 2000, from Cyprus Meteorological Department maps.
iii. Geology: the dominant underlying rock type, from Cyprus Geological Survey Department maps.
iv. Landscape diversity: apart from the scrub and pine forest habitat that was the focus for our surveys, some study sites were close to areas of cultivated farmland (cereals and/or tree crops). We used GIS analysis with the 2012 CORINE Land-cover (CLC) map for Cyprus as a layer to estimate the sum of the extent of all CLC categories corresponding to cultivation land use ^1^ within a 1 km radius around transect lines. We recorded landscape diversity as the proportion of scrub or pine forest within a 1 km radius around transect lines.

Table 2 (in appendix) shows the values for the above factors and the categorization for fire history for all study sites.

We excluded the potential confounding influence of three other factors through experimental design:

i. Disturbance from road traffic: no study site was closer than 1 km to a busy road. Mammides *et al*. (2014 and 2016) have shown that proximity to busy roads can have a negative impact on bird species richness in Cyprus.
ii. Hunting pressure: all study sites were within areas open to shooting in autumn and winter, so hunting pressure would be similar for all sites.
iii. Slope and Aspect: we ensured no study sites were on steep slopes.

### Analysis approach

#### Analysis of bird data from transects

As our focus was on breeding birds, records of mixed or single-species flocks of Linnet, European Goldfinch *Carduelis carduelis*, Greenfinch *Chloris chloris*, Serin *Serinus serinus*, and Corn Bunting *Emberiza calandra* encountered during counts in March were not included in the analysis, as they were likely to be winter visitors (Flint & Stewart 1992; Stylianou 2016, 2017, 2018). We also excluded early season records of species that do not breed in Cyprus, such as Blackcap *Sylvia atricapilla* and Chiffchaff *Phylloscopus collybita*. Also, and with the exception of aerial feeders such as hirundines, records of overflying birds were not included for analysis purposes as we could not be certain the birds were making use of the survey site habitat. They may have been just passing over en route between suitable habitat patches beyond the survey area.

We used the higher of the two counts for each species from the two site visits (early and late season). We chose this approach as the alternative of averaging from the two counts would have biased counts downwards for species with a distinct seasonality in their detectability and/or occurrence (such as European Roller, Sardinian Warbler *Sylvia melanocephala,* or Cyprus Warbler). Detectability of birds can be expected to vary between more open habitat types (phrygana and garrigue) and the more densely vegetated maquis and pine forest sites. We corrected for these differences in detectability by analysing our count data with Distance software (Thomas *et al*. 2010). We determined for each species separately the Effective Strip Width (ESW) values in phrygana, garrigue and maquis/pine forest habitat types. To enable this we categorized each of the 48 surveyed sites as phrygana, garrigue or maquis/forest using estimates of vegetation cover and density derived from vegetation surveys (see below), and based on the habitat classifications proposed by Tomaselli (1977):

i. Sites with > 10 % estimated cover in the > 2 m layer and an estimated 70 or more trees/tall bushes per hectare were classified as ‘Maquis/Forest’.
ii. Sites with < 10 % estimated cover in the > 2 m layer and estimated < 70 trees/tall bushes per hectare, but with > 10 % estimated cover in the 60 cm – 2m layer classified as ‘Garrigue’.
iii. Sites with < 5 % estimated cover in the > 2 m layer and estimated < 10 trees/tall bushes per hectare, and with < 10 % estimated cover in the 60 cm – 2m layer classified as ‘Phrygana’.

Using Distance analysis also allowed us to convert count data to estimated densities per hectare for each species. With ESW and transect length known, the effective area surveyed for each species for each site could be estimated, and thus a density estimate arrived at.

For each survey year, we calculated the following bird community metrics for each site:

i. Total abundance (sum of estimated mean densities/ha) and abundance of the subset of species of conservation priority (pooled).
ii. Overall species diversity, estimated using Simpson’s Index of Diversity:

Simpson’s Index of Diversity is 1 – D, where:

*D* = Σ(*n_i_*/*N*)^2^

and n = estimated density of species i in the sample

N = estimated density of all species in the sample

#### Analysis of vegetation data from quadrat and PCQM surveys

We averaged cover estimates for perennial vegetation in the three layers (0-60 cm, 60 cm-2 m, >2 m) and also for bare ground/rock and herbs/grasses, to arrive at estimates of percent cover for these parameters, and for individual perennial plant species, for each site. For perennials, we also calculated Simpson’s diversity index (as for birds, above, but using cover estimates instead of density estimates). We followed the analysis approach detailed by Mitchell (2015) to estimate density of trees/tall bushes per hectare from PCQM field data.

#### Statistical analysis approach

We built Generalized Linear Mixed Models (GLMMs) in lme4 in R3.5.1 (R Core Team 2018) to explore the relationships between response variables bird abundance, abundance of priority bird species and bird species diversity and grazing pressure and other explanatory variables. We built GLMMs in SPSS (IBM SPSS Statistics package v.20) to explore relationships between vegetation cover estimates and diversity of perennial vegetation and grazing pressure and other explanatory variables. We included the following additional predictor variables in our models: elevation, rainfall, geology (for vegetation models only), landscape diversity, fire history and its interaction with grazing pressure as fixed factors, and site and survey year (for the bird metrics only, as all vegetation variables were from 2017) as crossed random effects. For bird metrics, we also included diversity of perennial vegetation as a predictor variable. We transformed the dependent variables bird diversity and perennial vegetation diversity (power transformation ^6) and abundance (log transformation) to normalize their distribution, while the vegetation cover metrics were normally distributed. We also included the latitude and longitude of sites from the mid-point of transects in the model and tested for spatial autocorrelation in our models using Moran’s I tests. We found no evidence of spatial autocorrelation in any of our models. Best model fit was judged using the corrected Akaike’s Information Criterion (AICc) and was developed using backward selection from full models. Residuals were tested for normality using a combination of tests and plot examination, including skewness and kurtosis tests in Stata 11.2, Kdensity residual and QQnorm plots, and log(y) vs predicted/fitted values and fitted vs residual values correlations in R (Hawkins 2014).

## Results

In total, 9,245 individual birds of 58 species were recorded during 258 bird surveys carried out at 48 survey sites between mid-March and end of June of 2012, 2014 and 2016 (only 33 of the sites were covered in 2012). In addition, between January and March of 2017, vegetation surveys were carried out at the same 48 study sites. The total survey effort extended to 146 days over the four survey years, with a further 16 days of fieldwork in the preparatory stage (in late 2011), to locate survey sites and transect routes. Martin Hellicar carried out all this survey work.

Nineteen of the 58 bird species recorded were species of conservation priority (SPEC species and/or species listed in Annex I of the EU Birds Directive). Sixteen species of conservation concern were recorded in at least 10% of surveys, which we defined as regular occurrence (Table 3, in appendix). Among these priority birds were seven species for which Cyprus holds a significant proportion of the European breeding population (BirdLife International, 2004). These species were the two endemic breeding passerines, Cyprus Warbler and Cyprus Wheatear, plus Chukar Partridge *Alectoris chukar*, Black Francolin *Francolinus francolinus*, European Roller, Masked Shrike *Lanius nubicus*, and Cretzschmar’s Bunting.

The most commonly encountered species, recorded in at least 50% of sites, were Common Kestrel *Falco tinnunculus*, Chukar Partridge, Barn Swallow *Hirundo rustica*, Wood Pigeon *Columba palumbus*, Cyprus Wheatear, Sardinian Warbler, Cyprus Warbler, Great Tit *Parus major*, Magpie *Pica pica*, European Goldfinch, Greenfinch, and Cretzschmar’s Bunting. These twelve species are all common breeding birds in Cyprus (Flint & Stewart 1992, Stylianou 2016, 2017, 2018, Hellicar & Ieronymidou 2017).

Table 3 (in appendix) shows the density estimates for the sixteen regularly recorded bird species of conservation concern, plus the community metrics of overall abundance and species diversity. As we only carried out six bird surveys per site, it is unlikely we recorded all species present at sites, but our aim was to compare diversity between sites, rather than to arrive at an accurate measure of species richness.

Estimates of cover of perennial vegetation in the 0 – 60 cm layer varied from 8 – 73 % (average of 40.8 %). In the 60 cm – 2 m layer, the cover estimates varied from 0 – 46 % (average of 15.33 %), while in the > 2 m layer, cover varied from 0 – 35 % (average of 7.72 %). The PCQM estimate of tree/tall bush density varied from 0.07 to 1,136 per hectare (average of 93.27 per ha). Tables 1 and 2 (in appendix), show the vegetation cover and other estimates derived from the vegetation surveys for all sites.

The 48 study sites were classified for grazing pressure in the following way: eight ‘LOW’ grazing sites; nine grazing ‘RELEASE’ sites; twelve ‘MODERATE’ grazing sites; eight ‘INCREASED’ grazing sites and eleven ‘HIGH’ grazing sites. To gage the reliability of this categorization, we looked at how cover measures derived from vegetation surveys varied with grazing pressure categories. We found a pattern that suggested our categorization was reliable, with cover of perennial vegetation highest in low grazing and grazing release sites, and declining through the grazing sequence from moderate to increased to high (Figure 4). Woody vegetation would generally be expected to decrease with increasing pressure from grazing animals (Blondel & Aronson 1999, Allen 2001).

**Figure 4.**
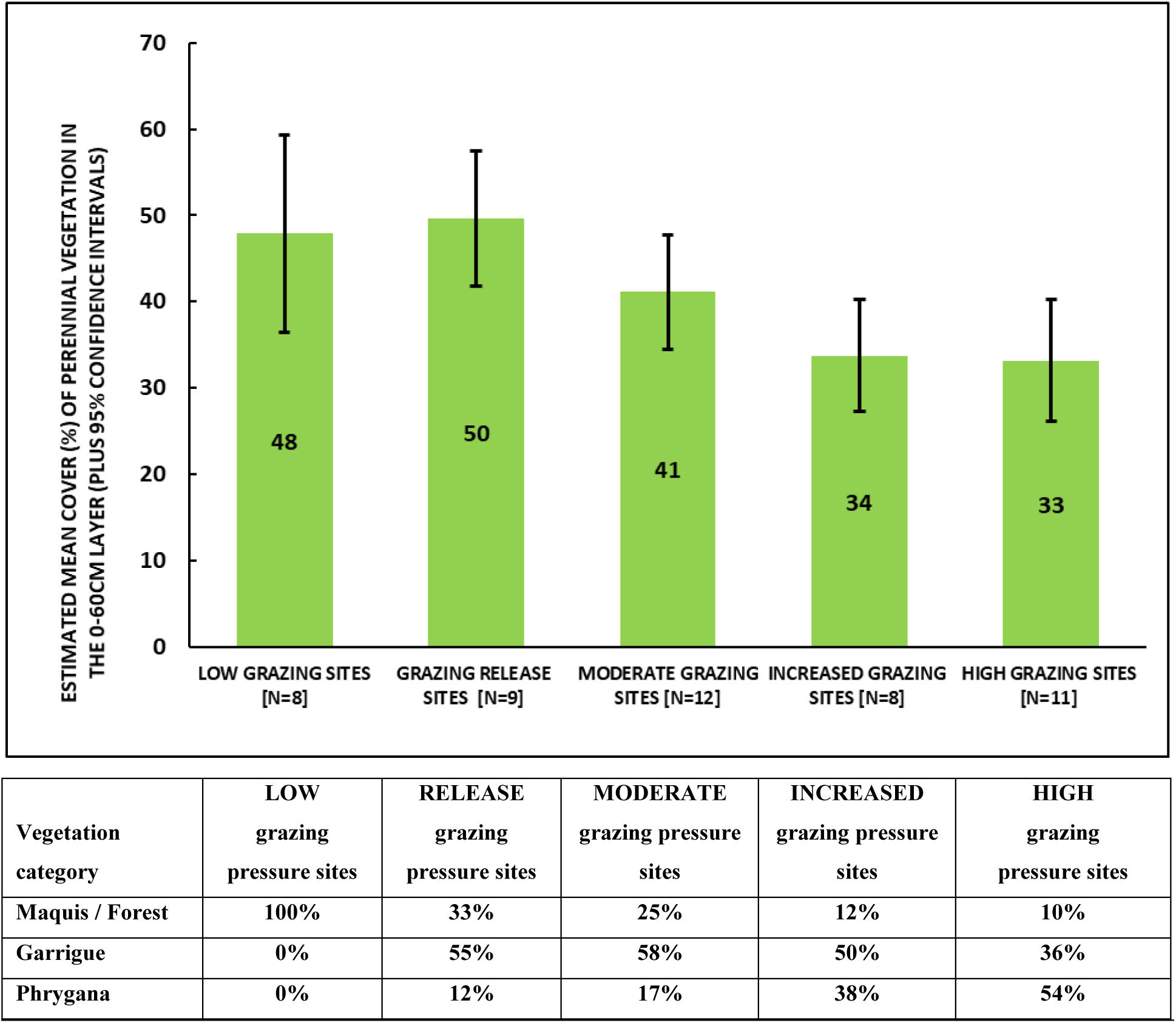
Mean perennial vegetation cover in the 0 - 60 cm layer in 2017 for sites under different grazing pressure regimes in Cyprus. The pattern is higher cover for sites with less grazing pressure (low grazing and release sites) than for sites with more grazing pressure (moderate, increased or high grazing sites). The table under the figure shows the proportion of sites classified as forest/maquis, garrigue and phrygana under each grazing pressure category.

In winter/early spring of 2017, only half of our study sites had sufficient ground cover of perennial and herbaceous vegetation to be at low risk from soil erosion, and if we looked at perennial plant cover alone, then only three of our 48 study sites met the cover threshold for preventing erosion set by Duran Zuazo & Rodriguez Pleguezuelo (2008). Even for sites classified as having the lowest grazing pressure, only one site (12.5 % of ‘low’ grazing sites) met the erosion-prevention threshold with perennial vegetation cover alone, while two (25 % of ‘low’ grazing sites) did not meet the threshold even when herbaceous plant cover was taken into account (see tables 1 and 2 in appendix).

Based on the habitat categorization used for the Distance analysis of bird count data (based on Tomaselli, 1977) 12 sites were classified as maquis/forest, 20 sites as garrigue and 16 sites as phrygana (Table 2 in appendix). The proportion of sites classified as maquis/forest under our system decreased with increasing grazing pressure, while, conversely, the proportion of phrygana sites increased. Sites classified as garrigue dominated the ‘mid-range’ grazing pressure categories (Figure 4). The most abundant (greatest mean cover) perennial plant species recorded in maquis and open forest sites were *Pinus brutia*, *Pistacia lentiscus*, *Cistus creticus*, *Cistus salviifolius*, *Juniperus phoenicea* and *Quercus alnifolia*. For garrigue sites, the most abundant perennials recorded were *Pistacia lentiscus*, *Genista fasselata fasselata, Cistus creticus, Cistus salviifolius*, and *Calycotome villosa*. For phrygana sites the most abundant perennials were the following: *Sarcopoterium spinosum*, *Thymus capitatus*, *Asphodelus ramosus*, *Cistus monspeliensis*, and *Noaea mucronata*.

Linear mixed models built for **overall bird abundance** showed there was no association between grazing pressure and bird numbers (Table 4 and Figure 7), but rather with fire history and diversity of perennial vegetation. There was a negative association between landscape diversity and overall abundance, showing that the presence of cultivated farmland (as well as scrub/forest) in the survey area was associated with higher bird numbers, but this pattern was borderline significant (t = −2.006, *P* = 0.051). The type of cultivation found within a 1 km radius of the survey transect consisted variously of cereals, vines and tree crops, mainly olives but also almonds, in a mosaic of small plots with remnants of semi-natural vegetation in between. The significant associations we found for overall bird abundance were with fire history and diversity of perennial vegetation. Bird abundance was lower in sites burnt sometime between just over a decade and five decades ago, i.e. sites categorized as ‘recovering’ (t = −4.218, *P* = 0.0001) or ‘recovered’ (t = −3.035, *P* = 0.004). Figure 3.5 shows the pattern of overall bird abundance for sites with different fire histories. ‘Unburnt’ sites, where there was no evidence of fire for the past half-century, show a pattern of higher bird numbers. Also, sites with higher diversity of perennial vegetation had higher bird abundances (t = 2.309, *P* = 0.026, Figure 6).

**Figure 5.**
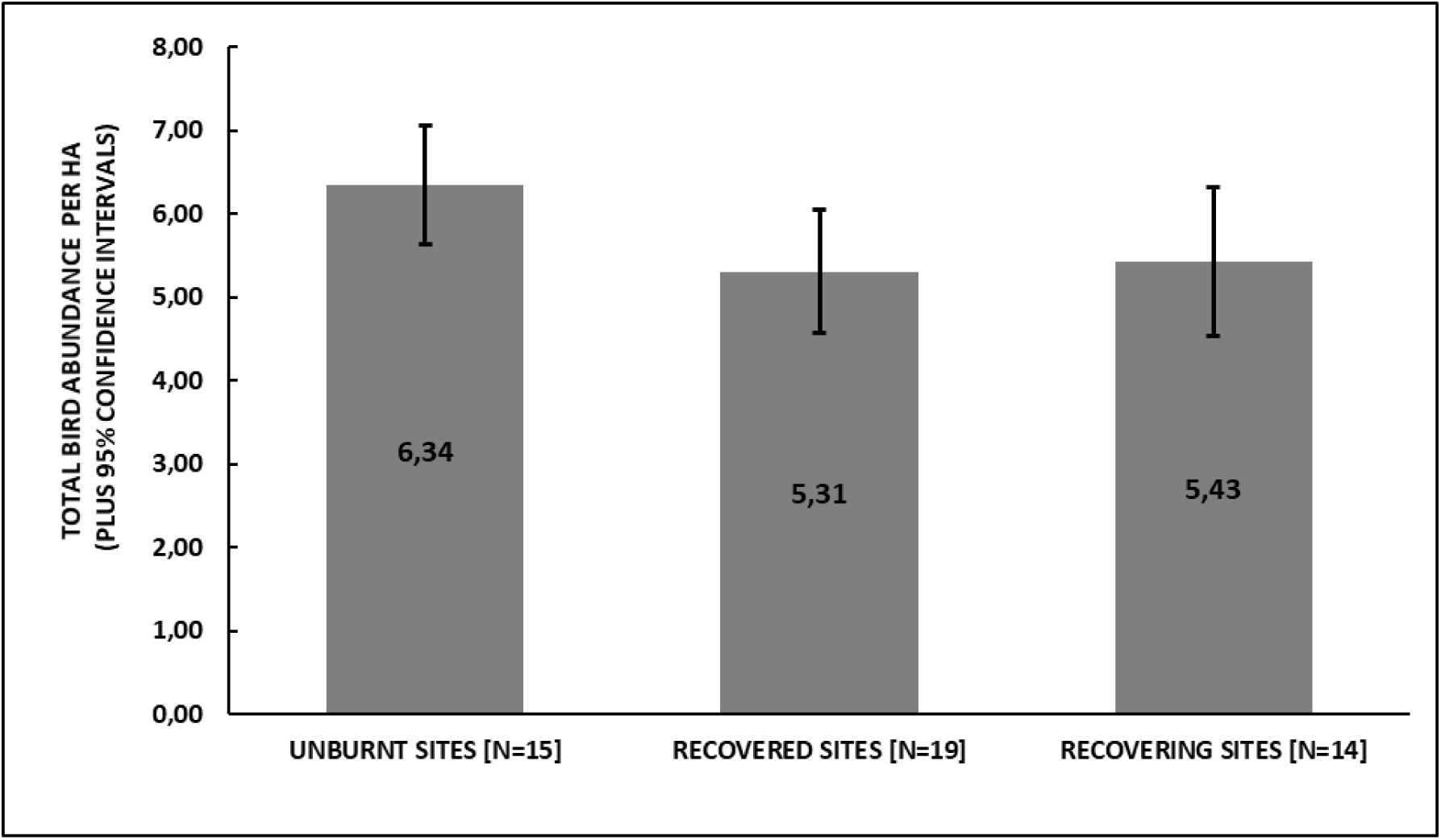
Overall bird abundance (sum of estimated mean species densities per ha, averaged from field survey data from 2012, 2014 and 2016) at study sites with different fire histories in Cyprus. Sites categorized as ‘unburnt’ (no evidence of fire for the past half-century) show a pattern for higher bird numbers than ‘recovered’ sites (last fire 20-55 years ago) or ‘recovering’ sites (last fire 12-20 years ago). The pattern is statistically significant at the *P* < 0.01 level.

**Figure 6.**
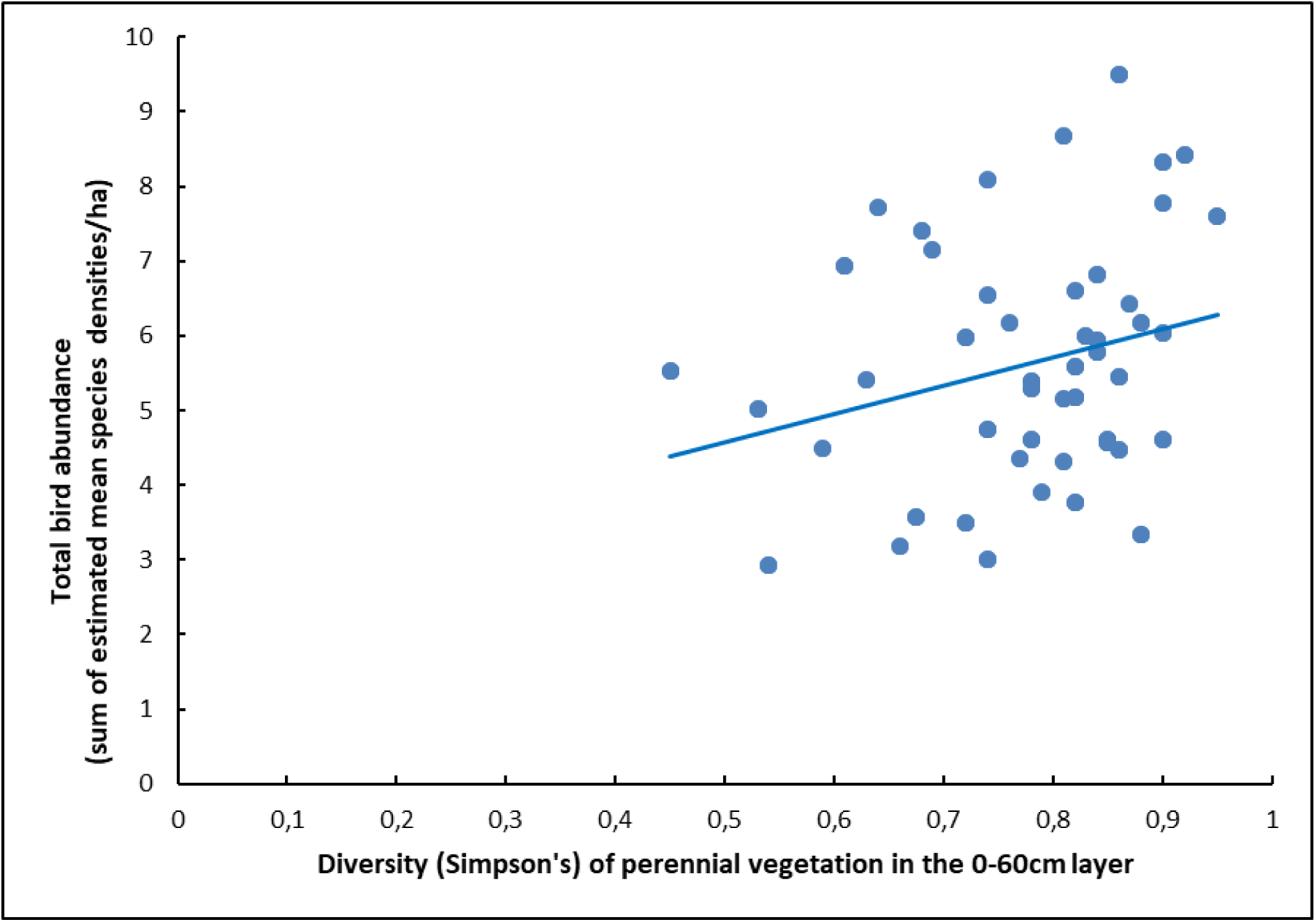
Association between overall bird abundance (sum of estimated mean species densities per ha, averaged from field survey data from 2012, 2014 and 2016) and diversity of perennial vegetation at study sites across Cyprus. The relationship is positive (as shown by line of best fit) with a higher diversity of perennial vegetation associated with higher bird abundance. The pattern is statistically significant (*P* = 0.026).

**Figure 7.**
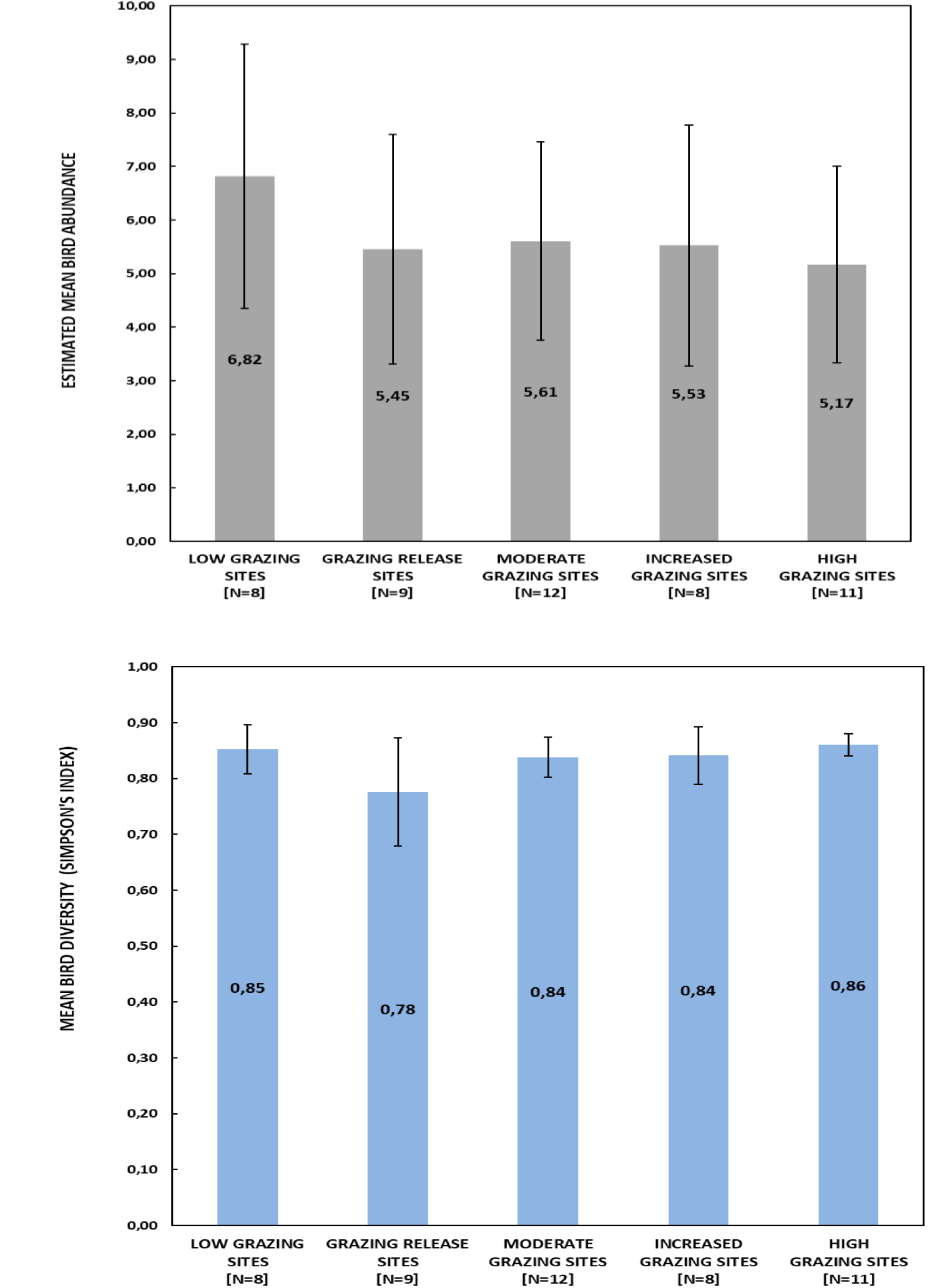
Patterns in bird abundance (per ha), top, and bird diversity (Simpson’s Index), bottom, averaged from field survey data from 2012, 2014 and 2016 (means with 95 % confidence intervals), at sites with different grazing pressures in Cyprus. There are no statistically significant differences between different site categories.

**Table 4.** Results for the full and best-supported GLMMs with the lowest AICc score under a Gaussian distribution model with overall bird abundance as response variable (highlighted in bold are variables included in the final model).

|  | **Estimate** | **St. Error** | **t value** | **P** |
| --- | --- | --- | --- | --- |
| **Intercept** | **0.818** | **0.198** | **4.131** | **<0.001** |
| **FIRE 1 ('recovered' sites)** | **-0.153** | **0.050** | **-3.035** | **0.004** |
| **FIRE 2 ('recovering' sites)** | **-0.243** | **0.057** | **-4.218** | **<0.001** |
| **Landscape diversity** | **-0.321** | **0.159** | **-2.006** | **0.051** |
| **Diversity perennial vegetation** | **0.470** | **0.204** | **2.309** | **0.026** |
| Grazing pressure 2 ('RELEASE') | -0.008 | 0.150 | 0.054 | 0.957 |
| Grazing pressure 3 ('MODERATE') | -0.159 | 0.085 | 1.883 | 0.069 |
| Grazing pressure 4 ('INCREASED') | 0.062 | 0.149 | 0.415 | 0.681 |
| Grazing pressure 5 ('HIGH') | -0.045 | 0.113 | 0.396 | 0.694 |
| Elevation | 0.121 | 0.059 | 2.040 | 0.049 |
| Rainfall | 0.001 | 0.004 | 0.143 | 0.887 |
| Grazing pressure 2 * Fire 1 | -0.037 | 0.211 | 0.176 | 0.861 |
| Grazing pressure 3 * Fire 1 | -0.025 | 0.182 | 0.139 | 0.890 |
| Grazing pressure 4 * Fire 1 | -0.280 | 0.218 | 1.286 | 0.209 |
| Grazing pressure 5 * Fire 1 | -0.102 | 0.191 | 0.532 | 0.598 |
| Grazing pressure 3 * Fire 2 | 0.0743 | 0.151 | 0.493 | 0.625 |
| Grazing pressure 4 * Fire 2 | -0.022 | 0.206 | 0.108 | 0.915 |
| *The AICc scores: best-supported model -79.764, full model -34.632* |  |  |  |  |

Linear mixed models built for **bird diversity** showed there was, again, no association with grazing pressure (Figure 7), but rather with landscape diversity and rainfall (Table 5). There was a negative association between landscape diversity and bird diversity, suggesting that the presence of cultivated farmland in the survey area was associated with greater bird diversity (t = −2.875, *P* = 0.006; Table 5). However, this association appears to be driven by two outlier data points, which undermines the validity of this finding. There was a clear positive association between rainfall (t = 4.740, *P* < 0.001) and bird diversity.

**Table 5.** Results for full and best-supported GLMMs with the lowest AICc score under a Gaussian distribution model with bird diversity as the response variable (highlighted in bold are variables included in the final model).

|  | **Estimate** | **St. Error** | **t value** | **P** |
| --- | --- | --- | --- | --- |
| **Intercept** | **-1.177** | **0.439** | **-2.681** | **0.010** |
| **Landscape diversity** | **-0.236** | **0.082** | **-2.875** | **0.006** |
| **Rainfall** | **0.810** | **0.171** | **4.740** | **<0.001** |
| Grazing pressure 2 ('RELEASE') | -0.104 | 0.080 | -1.295 | 0.204 |
| Grazing pressure 3 ('MODERATE') | -0.012 | 0.053 | -0.229 | 0.820 |
| Grazing pressure 4 ('INCREASED') | 0.059 | 0.076 | 0.774 | 0.445 |
| Grazing pressure 5 ('HIGH') | 0.030 | 0.065 | 0.467 | 0.643 |
| Fire 1 ('Recovered' sites) | 0.014 | 0.087 | 0.165 | 0.869 |
| Fire 2 ('Recovering' sites) | 0.027 | 0.064 | 0.431 | 0.669 |
| Elevation | 0.051 | 0.035 | 1.475 | 0.151 |
| Diversity perennial vegetation | 0.245 | 0.156 | 1.567 | 0.128 |
| Latitude | 0.152 | 0.079 | 1.917 | 0.065 |
| Longitude | -0.018 | 0.037 | 0.480 | 0.635 |
| Grazing pressure 2 * Fire 1 | 0.107 | 0.118 | 0.903 | 0.374 |
| Grazing pressure 3 * Fire 1 | 0.040 | 0.099 | 0.408 | 0.686 |
| Grazing pressure 4 * Fire 1 | -0.034 | 0.109 | -0.311 | 0.758 |
| Grazing pressure 5 * Fire 1 | 0.009 | 0.105 | 0.082 | 0.935 |
| Grazing pressure 3 * Fire 2 | -0.031 | 0.078 | -0.392 | 0.699 |
| Grazing pressure 4 * Fire 2 | -0.044 | 0.113 | -0.389 | 0.699 |
| *AICc scores: best-supported model -260.750, full model -171.982* |  |  |  |  |

Turning to the abundance of the subset of regularly recorded **bird species of conservation priority** (16 species, see Table 3 in appendix), we found a negative association with moderate grazing pressure (t = −2.426, *P* = 0.021; Table 6) and a positive association with elevation (t = 2.111, *P* = 0.043).

**Table 6.** Results for the full and best-supported GLMMs with the lowest AICc score under a Gaussian distribution model with abundance of 16 bird species of conservation priority as the response variable (highlighted in bold are variables included in the final model).

|  | **Estimate** | **St. Error** | **t value** | **P** |
| --- | --- | --- | --- | --- |
| **Intercept** | **73.963** | **50.488** | **1.465** | **0.153** |
| Landscape diversity | 0.606 | 2.199 | 0.276 | 0.785 |
| **Rainfall** | **-8.077** | **5.315** | **-1.520** | **0.139** |
| **Grazing pressure 2 ('RELEASE')** | **-1.345** | **1.759** | **-0.765** | **0.450** |
| **Grazing pressure 3 ('MODERATE')** | **-2.352** | **0.969** | **-2.426** | **0.021** |
| **Grazing pressure 4 ('INCREASED')** | **1.876** | **1.693** | **1.108** | **0.277** |
| **Grazing pressure 5 ('HIGH')** | **-1.505** | **1.357** | **-1.109** | **0.276** |
| **Fire 1 ('Recovered' sites)** | **-2.569** | **1.868** | **-1.375** | **0.179** |
| **Fire 2 ('Recovering' sites)** | **-0.843** | **1.382** | **-0.610** | **0.546** |
| **Elevation** | **1.622** | **0.768** | **2.111** | **0.043** |
| **Latitude** | **-1.552** | **1.269** | **-1.223** | **0.230** |
| Longitude | 0.721 | 0.840 | 0.858 | 0.398 |
| **Diversity perennial vegetation** | **4.183** | **3.253** | **1.286** | **0.209** |
| **Grazing pressure 2 \* Fire 1** | **3.084** | **2.601** | **1.186** | **0.245** |
| **Grazing pressure 3 \* Fire 1** | **2.640** | **2.109** | **1.251** | **0.221** |
| **Grazing pressure 4 \* Fire 1** | **-1.493** | **2.447** | **-0.610** | **0.547** |
| **Grazing pressure 5 \* Fire 1** | **1.620** | **2.993** | **0.705** | **0.487** |
| **Grazing pressure 3 \* Fire 2** | **-0.005** | **1.706** | **-0.003** | **0.998** |
| **Grazing pressure 4 \* Fire 2** | **-2.054** | **2.461** | **-0.835** | **0.411** |
| *AICc scores: best-supported model 458.897, full model 458.991* |  |  |  |  |

Linear mixed models built for **perennial vegetation cover metrics** (for the three vegetation layers: 0 – 60 cm, 60 cm – 2 m, > 2 m) showed there was an association with grazing pressure (see Figure 4 for the pattern for the 0 – 60 cm layer), and also with underlying geology and elevation. Results for the best-supported GLMMs showed the pattern was similar for all three vegetation layers (Table 7), with low grazing pressure sites having greater cover:

i. For the **0 – 60 cm vegetation layer**, there was a pattern of positive association between vegetation cover and both low grazing pressure (Wald χ^2^ = 5.839, *P* = 0.016) and grazing release sites (Wald χ^2^ = 4.391, *P* = 0.036). Cover in this layer was also positively associated with Gypsum-Sandstone geology (Wald χ^2^ = 12.460, *P* < 0.001) and negatively associated with Pillow lava geology (Wald χ^2^ = 4.043, *P* = 0.044), while landscape diversity (Wald χ^2^ = 4.153, *P* = 0.042) and elevation also had a positive association with cover (Wald χ^2^ = 10.015, *P* = 0.002).
ii. For the **60 cm – 2 m vegetation layer**, there was a pattern of positive association between vegetation cover and low grazing pressure (Wald χ^2^ = 9.440, *P* = 0.002).
iii. For the **> 2 m vegetation layer**, there was a pattern of positive association between vegetation cover and low grazing pressure (Wald χ^2^ = 12.362, *P* < 0.001). Cover in this layer was also positively associated with igneous geology (Wald χ^2^ = 4.060, *P* < 0.044).

**Table 7.** Results for the full and best-supported GLMMs with the lowest AICc score under a Gaussian distribution model with estimated cover of perennial vegetation in the (i) 0 – 60cm layer,(ii) 60 cm – 2 m layer and (iii) >2 m layer, as the response variable (highlighted in bold are variables included in the final model).

| (i) | Estimate | St. Error | Wald $\chi^2$ | P |
| --- | --- | --- | --- | --- |
| **Intercept** | **49.392** | **18.409** | **7.199** | **0.007** |
| **Grazing pressure 1 ('LOW')** | **21.406** | **8.858** | **5.839** | **0.016** |
| **Grazing pressure 2 ('RELEASE')** | **13.619** | **6.499** | **4.391** | **0.036** |
| **Grazing pressure 3 'MODERATE')** | **-0.875** | **6.027** | **0.021** | **0.885** |
| **Grazing pressure 4 ('INCREASED')** | **-8.575** | **6.645** | **1.665** | **0.197** |
| Rainfall | -0.083 | 0.046 | 3.189 | 0.074 |
| **Elevation** | **0.038** | **0.012** | **10.015** | **0.002** |
| **Mamonia geology** | **3.374** | **7.651** | **0.194** | **0.659** |
| **Clay geology** | **-8.141** | **7.971** | **1.043** | **0.307** |
| **Gypsum-Sandstone geology** | **28.008** | **7.937** | **12.460** | **<0.001** |
| **Chalk geology** | **-3.747** | **6.583** | **0.279** | **0.598** |
| **Pillow lava geology** | **24.392** | **12.130** | **4.043** | **0.044** |
| **Igneous geology** | **14.853** | **8.301** | **3.202** | **0.074** |
| Fire 0 ('unburnt sites') | -1.010 | 7.836 | 0.017 | 0.897 |
| Fire 1 ('Recovered' sites) | -11.323 | 6.414 | 3.116 | 0.078 |
| **Landscape diversity** | **28.194** | **13.834** | **4.153** | **0.042** |
| Grazing pressure 1 * Fire 0 | -24.137 | 12.649 | 3.641 | 0.056 |
| Grazing pressure 3 * Fire 0 | 0.203 | 9.813 | 0.000 | 0.984 |
| Grazing pressure 4 * Fire 0 | -18.010 | 12.721 | 2.004 | 0.157 |
| Grazing pressure 2 * Fire 1 | 3.438 | 9.491 | 0.131 | 0.717 |
| Grazing pressure 3 * Fire 1 | 8.608 | 8.311 | 1.073 | 0.300 |
| Grazing pressure 4 * Fire 1 | 16.832 | 8.919 | 3.561 | 0.059 |
| *AICc scores: best-supported model 424.431, full model 603.309* |  |  |  |  |
| (ii) | Estimate | St. Error | Wald $\chi^2$ | P |
| **Intercept** | **4.280** | **17.877** | **0.057** | **0.811** |
| **Grazing pressure 1 ('LOW')** | **26.430** | **8.602** | **9.440** | **0.002** |
| **Grazing pressure 2 ('RELEASE')** | **4.316** | **6.311** | **0.468** | **0.494** |
| **Grazing pressure 3 'MODERATE')** | **-1.196** | **5.853** | **0.042** | **0.838** |
| **Grazing pressure 4 ('INCREASED')** | **-3.056** | **6.453** | **0.224** | **0.636** |
| Rainfall | -0.004 | 0.449 | 0.009 | 0.924 |
| Elevation | 0.004 | 0.012 | 0.138 | 0.710 |
| **Mamonia geology** | **0.866** | **7.429** | **0.014** | **0.907** |
| **Clay geology** | **-5.471** | **7.741** | **0.500** | **0.480** |
| **Gypsum-Sandstone geology** | **12.224** | **7.705** | **2.517** | **0.113** |
| **Chalk geology** | **2.427** | **6.392** | **0.144** | **0.704** |
| **Pillow lava geology** | **-9.068** | **11.779** | **0.593** | **0.441** |
| **Igneous geology** | **-2.400** | **8.061** | **0.089** | **0.766** |
| Fire 0 ('unburnt sites') | 2.788 | 7.609 | 0.134 | 0.714 |
| Fire 1 ('Recovered' sites) | -6.297 | 6.229 | 1.022 | 0.312 |
| Landscape diversity | 9.767 | 13.434 | 0.528 | 0.467 |
| Grazing pressure 1 * Fire 0 | -21.044 | 12.283 | 2.935 | 0.087 |
| Grazing pressure 3 * Fire 0 | 5.483 | 9.529 | 0.331 | 0.565 |
| Grazing pressure 4 * Fire 0 | 3.993 | 12.353 | 0.104 | 0.747 |
| Grazing pressure 2 * Fire 1 | 7.604 | 9.217 | 0.681 | 0.409 |
| Grazing pressure 3 * Fire 1 | 10.119 | 8.070 | 1.572 | 0.210 |
| Grazing pressure 4 * Fire 1 | 4.327 | 8.661 | 0.250 | 0.617 |
| *AICc scores: best-supported model 421.614, full model 556.908* |  |  |  |  |
| **(iii)** | **Estimate** | **St. Error** | **Wald $\chi^2$** | **P** |
| **Intercept** | **-6.407** | **8.070** | **0.630** | **0.427** |
| **Grazing pressure 1 ('LOW')** | **13.654** | **3.883** | **12.362** | **<0.001** |
| **Grazing pressure 2 ('RELEASE')** | **-1.938** | **2.849** | **0.463** | **0.496** |
| **Grazing pressure 3 'MODERATE')** | **-2.685** | **2.642** | **1.033** | **0.309** |
| **Grazing pressure 4 ('INCREASED')** | **-0.407** | **2.913** | **0.019** | **0.889** |
| Rainfall | 0.016 | 0.020 | 0.589 | 0.443 |
| Elevation | 0.001 | 0.005 | 0.014 | 0.904 |
| **Mamonia geology** | **0.270** | **3.354** | **0.006** | **0.936** |
| **Clay geology** | **-2.894** | **3.494** | **0.868** | **0.408** |
| **Gypsum-Sandstone geology** | **4.382** | **3.475** | **1.587** | **0.208** |
| **Chalk geology** | **1.310** | **2.886** | **0.206** | **0.650** |
| **Pillow lava geology** | **-0.859** | **5.318** | **0.026** | **0.872** |
| **Igneous geology** | **7.332** | **3.639** | **4.060** | **0.044** |
| **Fire 0 ('unburnt sites')** | **3.611** | **3.435** | **1.105** | **0.293** |
| **Fire 1 ('Recovered' sites)** | **-1.319** | **2.812** | **0.220** | **0.639** |
| Landscape diversity | 1.918 | 6.065 | 0.100 | 0.752 |
| Grazing pressure 1 * Fire 0 | -5.745 | 5.545 | 1.073 | 0.300 |
| Grazing pressure 3 * Fire 0 | 3.792 | 4.302 | 0.777 | 0.378 |
| Grazing pressure 4 * Fire 0 | 2.297 | 5.577 | 0.170 | 0.680 |
| Grazing pressure 2 * Fire 1 | 3.061 | 4.161 | 0.541 | 0.462 |
| Grazing pressure 3 * Fire 1 | 7.108 | 3.643 | 3.806 | 0.051 |
| Grazing pressure 4 * Fire 1 | 0.624 | 3.910 | 0.025 | 0.873 |
| *AICc scores: best-supported model 345.264, full model 452.909* |  |  |  |  |

Turning to **perennial plant diversity** (Simpson’s index), the best-supported Linear mixed model (Table 8) showed a positive association with low grazing pressure sites compared to high grazing pressure sites and a pattern (non-significant) of positive association with sites where grazing was moderate or had increased in recent years, again compared to high grazing pressure sites. There was also a positive association of plant diversity with rainfall.

**Table 8.** Results for the full and best-supported GLMMs with the lowest AICc score under a Gaussian distribution model with diversity of perennial vegetation as the response variable (highlighted in bold are variables included in the final model).

|  | **Estimate** | **St. Error** | **Wald $\chi^2$** | **P** |
| --- | --- | --- | --- | --- |
| **Intercept** | **0.807** | **0.206** | **15.288** | **< 0.001** |
| **Grazing pressure 1 ('LOW')** | **0.322** | **0.118** | **7.483** | **0.006** |
| **Grazing pressure 2 ('RELEASE')** | **-0.030** | **0.083** | **0.132** | **0.716** |
| **Grazing pressure 3 ('MODERATE')** | **0.056** | **0.081** | **0.481** | **0.488** |
| **Grazing pressure 4 ('INCREASED')** | **0.100** | **0.088** | **1.298** | **0.255** |
| **Rainfall** | **0.001** | **0.0004** | **4.957** | **0.026** |
| Mamonia geology | 0.206 | 0.089 | 5.404 | 0.020 |
| Clay geology | 0.164 | 0.102 | 2.596 | 0.107 |
| Gypsum-Sandstone geology | -0.012 | 0.105 | 0.012 | 0.912 |
| Chalk geology | 0.086 | 0.077 | 1.258 | 0.262 |
| Pillow lava geology | 0.099 | 0.124 | 0.633 | 0.426 |
| Igneous geology | 0.052 | 0.080 | 0.421 | 0.516 |
| **Fire 0 ('unburnt sites')** | **0.139** | **0.103** | **1.829** | **0.176** |
| **Fire 1 ('Recovered' sites)** | **0.060** | **0.086** | **0.489** | **0.484** |
| **Grazing pressure 1 \* Fire 0** | **0.293** | **0.168** | **3.063** | **0.080** |
| **Grazing pressure 2 \* Fire 1** | **- 0.012** | **0.119** | **0.009** | **0.923** |
| **Grazing pressure 3 \* Fire 0** | **- 0.183** | **0.131** | **1.962** | **0.161** |
| **Grazing pressure 3 \* Fire 1** | **0.064** | **0.110** | **0.336** | **0.562** |
| *AICc scores: best-supported model -4.063, full model 127.078* |  |  |  |  |

## Discussion

Our results showed a clear pattern of association between grazing pressure and vegetation, but patterns linking grazing pressure to the bird community were not as clear-cut. This despite the fact that our island-wide study achieved a balanced field experiment design; with similar (and adequate) sample size in each of the five categories we defined to capture current and past grazing pressure: low, release, moderate, increase and high.

The influence of grazing on birds is likely to be through the role of sheep and goats as ‘ecosystem engineers’, generating a more patchy and heterogeneous habitat (Shachak *et al*. 2008, Arga & Ne’eman 2009, Grove & Rackham 2001), rather than through direct interaction of livestock with birds. Livestock might disturb ground-nesting birds such as Chukar Partridge and Black Francolin (Snow & Perrins 1998), and some bird species, such as Cattle Egret *Bubulcus ibis*, wagtails *Motacilla spp* and some members of the crow family Corvidae are known to follow grazing animals to feed on invertebrates that livestock disturb (Snow & Perrins 1998). Egrets and wagtails were not recorded at our study sites, and though we did observe Jackdaws *Corvus monedula* associating directly with sheep and goat herds on four occasions, this was the only direct interaction between birds and livestock we recorded.

We identified a clear pattern for grazing pressure influencing the structure and composition of perennial vegetation. As would be expected, we found an overall pattern linking increasing grazing pressure to decreasing vegetation cover. Both diversity and cover (in all vegetation layers) of perennials were positively associated with sites categorised as subject to low grazing pressure, both currently and in the past. In the case of ground-level perennial vegetation (in the 0 – 60 cm layer), there was also a positive association with sites where grazing pressure was heavier in the past, but had been reduced in more recent years (‘release’ sites).

Reduction of overall perennial vegetation cover due to grazing pressure is a pattern recorded in many other parts of the Mediterranean (Blondel & Aronson, 1999, Grove & Rackham 2001, Allen 2001, Papanastasis, Kyriakakis & Kazakis 2002, Blondel *et al*. 2010). The increased cover in ‘release’ sites can be interpreted as a response to a reduction in browsing impact. The pattern for this being seen only in the 0 – 60 cm vegetation layer could be due to this lifting of browsing pressure being relatively recent in Cyprus. This would mean there has not yet been sufficient time for the cover increase resulting from the release of grazing to materialise in higher vegetation layers. The positive association we found for cover in the >2 m layer with igneous geology is explained by pine wood habitat sites, with their greater tree cover, being largely restricted to the igneous rocks of the Troodos Mountains.

Our finding that sites with the lowest levels of grazing pressure had higher perennial plant diversity than overgrazed sites appears to be inconsistent with what has been shown for other parts of the Mediterranean, where continuous low-level grazing pressure was found to generally increase floristic diversity (Naveh & Whittaker 1979, Montalvo *et al*. 1993, Noy-Meir 1998, Papanastasis 1998, Peco *et al*. 1998, Grove & Rackham 2001, Papanastasis, Kyriakakis & Kazakis 2002, Shachak *et al*. 2008, Arga & Ne’eman 2009, Blondel *et al*. 2010). However, it is worth noting that the studies in other parts of the Mediterranean looked at herbaceous as well as woody vegetation, something we did not do. Diversity of perennial vegetation was also positively associated with rainfall, with wetter areas having a greater variety of woody plants.

Looking for direct associations between birds and grazing pressure, the only significant result was a negative association between the abundance of the sub-set of bird species of conservation priority and sites subject to moderate grazing pressure, both currently and in the past. Overall bird abundance was not clearly associated with grazing pressure, and nor was overall bird diversity.

We did however, find a positive association between perennial plant diversity and overall bird abundance, which points to a link between grazing pressure and breeding bird numbers, as high diversity of perennial plants was clearly associated with low grazing pressure. But a similar pattern was not seen for bird diversity or abundance of priority bird species, which were not significantly associated with vegetation parameters.

Of the other factors associated with bird community metrics, only one, fire, was largely anthropogenic. For overall bird abundance, our models linked higher breeding bird abundance with sites where we identified no evidence of perturbation by fire for at least 50 years before our study began.

Rainfall was clearly and positively associated with breeding bird diversity, while in the case of the 16 bird species of priority conservation concern regularly recorded in our study, there was also a positive association with elevation. Cyprus is a generally arid island, and in such arid environments, bird species richness has been shown to increase with rainfall (Seymour *et al*. 2015). This is a likely explanation for the pattern we found for bird diversity. Among the priority bird species were the endemic Cyprus Wheatear and Cyprus Warbler and five other species for which Cyprus holds a significant proportion of the European breeding population: Chukar Partridge, Black Francolin, European Roller, Masked Shrike, and Cretzschmar’s Bunting. The positive link of this species sub-set with elevation is probably explained to a large extent by the greater prevalence of forest habitat at higher altitude. The Cyprus Wheatear, Masked Shrike, and Cretzschmar’s Bunting are species that tend to reach higher abundance levels in forest habitat in Cyprus (Flint & Stewart 1992, Stylianou 2016, 2017, 2018, Hellicar *et al*. 2014) and we recorded all three species in higher numbers at higher altitude sites.

We aimed to test whether the intermediate disturbance hypothesis (Blondel & Aronson 1999 and Allen 2001, Blondel *et al*. 2010) would hold true for Cyprus when it comes to grazing by sheep and goats, as well as for other anthropogenic disturbance factors, such as fire. For this to be supported, a greater diversity of breeding birds and of plants would need to be found where habitats have been subject to continuous low-level perturbation over a long period of time. We would also expect lower diversity of birds and of plants at sites where disturbance has recently ceased, has been intensive over time or has occurred only very rarely or not at all. The IDH postulates that this would be true in particular for local scale or α-diversity, which was what we focused on in this study.

What we found for grazing, was that sites we classified as low disturbance were the ones associated with a greater diversity of perennial vegetation. Our ‘low’ grazing category sites – where grazing was often absent and at least very limited - also had greater cover of perennial vegetation. These high plant diversity sites were in turn associated with greater abundance of breeding birds. However, breeding bird diversity was not associated with grazing either directly or indirectly. Of note was the negative association between abundance of the sub-set of bird species of priority conservation concern and sites categorized as having been under a long-term regime of moderate grazing pressure.

As diversity is the key metric in relation to the IDH, our findings for grazing seem to contradict the IDH, in that regular low-level perturbation was not found to be linked to greater biodiversity (at least not for breeding birds or perennial plants). Greater local-level diversity of woody plants was instead associated with sites within designated State Forest areas. In these areas, grazing by goats and sheep is generally banned and has been largely absent – or at least infrequent and very low-level – for over a century (with designation of State Forests beginning as long ago as 1879).

Our finding relating to fire, another key disturbance factor for Mediterranean systems, also runs counter to the IDH. We found no evidence of the interaction between grazing and fire having a significant impact on birds or plants. We did however find that breeding bird abundance was greater at sites that had not been perturbed by fire for at least half a century before our study began. Caveats here are that these results relate to bird abundance and not diversity, and that our experimental design aimed at excluding the potentially confounding influence of fire, so we did not sample any areas burnt after 2000. The latter meant the impact of recent burns and their interaction with grazing were not within the scope of the current study.

For Cyprus therefore, our findings for grazing and fire suggest that what is beneficial for biodiversity is the lowest levels of perturbation we defined, which were equivalent to a general absence of disturbance. But it is worth noting that our approach for defining past and current grazing pressure at our study sites enabled us to arrive only at a categorization for relative grazing pressure, rather than a quantified, continuous measure of site use by animals. Based on vegetation cover estimates, at least half of our study sites overall - and 25 % of low grazing pressure sites - could be classified as overgrazed, as they fell below the 65-70 % vegetation cover threshold for prevention of soil erosion (Duran Zuazo and Rodriguez Pleguezuelo 2008). If we consider only perennial vegetation, and not the seasonal herbaceous plants, then only three of our 48 sites met the erosion-prevention threshold of 65-70 % plant cover. This suggests our approach to grazing pressure categorization may have underestimated actual grazing pressure. Another source of underestimation derives from the fact that the number of goats and sheep currently in pens is often higher than the number licensed by the Veterinary Service. Nonetheless, the approach we adopted was based on best available data. While it did not allow us to arrive at a reliable quantification of actual numbers of grazing animal per unit area, it did amount to a consistent relative measure for comparing grazing pressure (past and present) at different study sites.

Our categorization of sites for fire history was also relative and to an extent approximate, because definite records of last burns were not available for the period before 2000. Both these factors introduce a level of uncertainty and they should be borne in mind when interpreting our findings. Different patterns might have emerged from an analysis based on more precise data on the numbers of grazing animals visiting sites (and the frequency of their visits) and on date-of-last-burn.

While noting these methodological caveats, the pattern we identified for the influence of fire and grazing on birds and vegetation suggest that it is not a regime of continuous low-level ecosystem disturbance, but rather the absence of perturbation – or at least only very minimal perturbation – that is beneficial for biodiversity in scrub and open woodland habitats in Cyprus. This finding suggests the best approach for biodiversity management in scrub and open woodland habitats in Cyprus is to aim to avoid fire and to keep grazing to a minimum. However, we would not support the adoption of a ‘no grazing’ approach for scrub or forest management. There are two reasons for this. Firstly, we may have underestimated grazing pressure in our study, as noted above, which would mean our ‘low’ grazing pressure sites may in fact have been subject to regular grazing. Secondly, absence of grazing would significantly increase the risk of fire – with, as we have shown, negative impact on biodiversity - due to accumulation of combustible plant material in the system (Blumler 1996, Grove & Rackham 2001, Allen 2001, Papanastasis, Kyriakakis & Kazakis 2002, Blondel *et al*. 2010).

## APPENDICES

**Table 3.1.** Grazing characteristics and grazing pressure categorization for survey sites (See text for explanations of how these parameters were estimated)

| Site identity |  | Grazing parameters and grazing pressure categorisation |  |  |  |  |  |  |  |  |  |  |
| --- | --- | --- | --- | --- | --- | --- | --- | --- | --- | --- | --- | --- |
| Study site name | Study area | Grazing animals within site | Grazing animals within 1 km radius of site | Grazing animals within 3 km radius of site | Estimated total grazing animal pressure on site | CURRENT GRAZING PRESSURE CATEGORISATION | Site categorisation under 1935 Goat Law | HISTORICAL GRAZING PRESSURE CATEGORISATION | OVERALL GRAZING PRESSURE CATEGORISATION | Average bush compactness measure | Sum of grazing animal tracks recorded on site | Sum of grazing animal droppings recorded on site |
| Akamas1 | Akamas | 0 | 185 | 2191 | 623,2 | HIGH | STATE FOREST | HIGH | HIGH | 21 | 14 | 175 |
| Akamas2 | Akamas | 0 | 0 | 1157 | 231,4 | MODERATE | STATE FOREST | MODERATE | MODERATE | 13 | 7 | 20 |
| Akamas3 | Akamas | 0 | 0 | 400 | 80 | MODERATE | STATE FOREST | MODERATE | MODERATE | 47,5 | 5 | 25 |
| Akamas4 | Akamas | 0 | 34 | 3261 | 686,2 | HIGH | PRESCRIBED VILLAGE AREA | MODERATE | INCREASE | 21 | 11 | 120 |
| Akamas5 | Akamas | 0 | 100 | 0 | 100 | MODERATE | STATE FOREST | MODERATE | MODERATE | 15 | 8 | 150 |
| Akamas6 | Akamas | 57 | 128 | 2134 | 668,8 | HIGH | NON-PRESCRIBED VILLAGE AREA | HIGH | HIGH | 32,5 | 12 | 155 |
| Kapras4 | Karpasia | 0 | 0 | 150 | 30 | LOW | STATE FOREST | LOW | LOW | 52 | 3 | 5 |
| Karpas1 | Karpasia | 0 | 200 | 100 | 220 | MODERATE | STATE FOREST | MODERATE | LOW | 53 | 9 | 150 |
| Karpas2 | Karpasia | 200 | 0 | 0 | 400 | HIGH | STATE FOREST | MODERATE | INCREASE | 71 | 4 | 261 |
| Karpas6 | Karpasia | 0 | 0 | 0 | 0 | LOW | STATE FOREST | LOW | LOW | 71 | 0 | 0 |
| Karpas7 | Karpasia | 60 | 70 | 250 | 240 | MODERATE | PRESCRIBED VILLAGE AREA | MODERATE | MODERATE | 47 | 4 | 0 |
| Karpas NScrub | Karpasia | 0 | 0 | 320 | 64 | MODERATE | NON-PRESCRIBED VILLAGE AREA | HIGH | DECREASE | 18,5 | 4 | 25 |
| Dali | Larnaca | 0 | 378 | 1927 | 763,4 | HIGH | PRESCRIBED VILLAGE AREA | MODERATE | INCREASE | 28 | 9 | 20 |
| Kalavassos 1 | Larnaca | 0 | 0 | 0 | 0 | LOW | PRESCRIBED VILLAGE AREA | MODERATE | DECREASE | 32 | 2 | 0 |
| Kalavassos 3 | Larnaca | 444 | 0 | 2451 | 1378,2 | HIGH | PRESCRIBED VILLAGE AREA | MODERATE | INCREASE | 31 | 17 | 110 |
| Lefkara 1 | Larnaca | 0 | 0 | 180 | 36 | LOW | PRESCRIBED VILLAGE AREA | MODERATE | DECREASE | 33 | 2 | 0 |
| Lefkara 2 | Larnaca | 180 | 0 | 956 | 551,2 | HIGH | PRESCRIBED VILLAGE AREA | MODERATE | INCREASE | 40 | 15 | 150 |
| Lefkara 3 | Larnaca | 0 | 0 | 0 | 0 | LOW | PRESCRIBED VILLAGE AREA | MODERATE | DECREASE | 58 | 2 | 0 |
| Maroni 2 | Larnaca | 0 | 0 | 1357 | 271,4 | MODERATE | PRESCRIBED VILLAGE AREA | MODERATE | MODERATE | 48 | 0 | 0 |
| Maroni 3 | Larnaca | 0 | 0 | 321 | 64,2 | MODERATE | PRESCRIBED VILLAGE AREA | MODERATE | MODERATE | 57 | 2 | 0 |
| Maroni 4 | Larnaca | 0 | 904 | 453 | 994,6 | HIGH | PRESCRIBED VILLAGE AREA | MODERATE | INCREASE | 71 | 5 | 45 |
| Stazousa 1 | Larnaca | 0 | 0 | 1658 | 331,6 | MODERATE | NON-PRESCRIBED VILLAGE AREA | HIGH | DECREASE | 87 | 10 | 0 |
| Stazousa 2 | Larnaca | 0 | 227 | 892 | 405,4 | HIGH | NON-PRESCRIBED VILLAGE AREA | HIGH | HIGH | 64 | 7 | 60 |
| Stazousa 3 | Larnaca | 631 | 408 | 3053 | 2280,6 | HIGH | NON-PRESCRIBED VILLAGE AREA | HIGH | HIGH | 25 | 17 | 125 |
| Episkopi | Limassol | 0 | 194 | 0 | 194 | MODERATE | STATE FOREST | LOW | LOW | 48 | 6 | 260 |
| Monagri1 | Limassol | 0 | 0 | 1035 | 207 | MODERATE | PRESCRIBED VILLAGE AREA | MODERATE | MODERATE | 47 | 5 | 60 |
| Paramali1 | Limassol | 0 | 512 | 4263 | 1364,6 | HIGH | NON-PRESCRIBED VILLAGE AREA | HIGH | HIGH | 83 | 9 | 70 |
| Paramali2 | Limassol | 0 | 0 | 4417 | 883,4 | MODERATE | PRESCRIBED VILLAGE AREA | MODERATE | MODERATE | 17 | 4 | 10 |
| Petra Romiou1 | Limassol | 486 | 884 | 970 | 2050 | HIGH | NON-PRESCRIBED VILLAGE AREA | HIGH | HIGH | 35 | 13 | 245 |
| Petra Romiou2 | Limassol | 0 | 442 | 1190 | 680 | HIGH | STATE FOREST | MODERATE | INCREASE | 42 | 7 | 15 |
| Analyontas 2 | Nicosia | 0 | 0 | 2147 | 429,4 | HIGH | NON-PRESCRIBED VILLAGE AREA | HIGH | HIGH | 29 | 6 | 0 |
| Analyontas 3 | Nicosia | 0 | 181 | 903 | 361,6 | MODERATE | NON-PRESCRIBED VILLAGE AREA | HIGH | DECREASE | 30 | 5 | 0 |
| Delikipos 1 | Nicosia | 0 | 0 | 170 | 34 | LOW | PRESCRIBED VILLAGE AREA | MODERATE | DECREASE | 77 | 5 | 0 |
| Delikipos 2 | Nicosia | 0 | 0 | 0 | 0 | LOW | PRESCRIBED VILLAGE AREA | MODERATE | DECREASE | 60 | 0 | 0 |
| Dyo Moutes | Nicosia | 0 | 0 | 0 | 0 | LOW | STATE FOREST | LOW | LOW | 76 | 0 | 0 |
| Kapedes 1 | Nicosia | 0 | 0 | 462 | 92,4 | MODERATE | PRESCRIBED VILLAGE AREA | MODERATE | MODERATE | 48 | 2 | 0 |
| Kapedes 2 | Nicosia | 0 | 0 | 643 | 128,6 | MODERATE | PRESCRIBED VILLAGE AREA | MODERATE | MODERATE | 19 | 1 | 0 |
| Kionia | Nicosia | 0 | 0 | 0 | 0 | LOW | STATE FOREST | LOW | LOW | 39 | 1 | 0 |
| Machairas 1 | Nicosia | 0 | 54 | 392 | 132,4 | MODERATE | STATE FOREST | LOW | LOW | 70 | 17 | 50 |
| Machairas 2 | Nicosia | 0 | 0 | 54 | 10,8 | LOW | STATE FOREST | LOW | LOW | 36 | 0 | 0 |
| Xilias 1 | Nicosia | 0 | 396 | 174 | 430,8 | MODERATE | NON-PRESCRIBED VILLAGE AREA | HIGH | DECREASE | 60 | 3 | 0 |
| Xilias 2 | Nicosia | 0 | 396 | 112 | 418,4 | HIGH | NON-PRESCRIBED VILLAGE AREA | HIGH | HIGH | 30 | 4 | 15 |
| Diarizos1 | Xeros/Diarizos | 0 | 700 | 2447 | 1189,4 | HIGH | NON-PRESCRIBED VILLAGE AREA | HIGH | HIGH | 44 | 18 | 25 |
| Diarizos2 | Xeros/Diarizos | 0 | 1723 | 2190 | 2161 | HIGH | NON-PRESCRIBED VILLAGE AREA | HIGH | HIGH | 60 | 5 | 70 |
| Xeros 5 | Xeros/Diarizos | 0 | 0 | 1479 | 295,8 | MODERATE | PRESCRIBED VILLAGE AREA | MODERATE | MODERATE | 74 | 5 | 0 |
| Xeros1 | Xeros/Diarizos | 0 | 0 | 1094 | 218,8 | MODERATE | mixed prescribed and not | MODERATE | MODERATE | 67 | 3 | 0 |
| Xeros2 | Xeros/Diarizos | 0 | 390 | 1297 | 649,4 | HIGH | NON-PRESCRIBED VILLAGE AREA | HIGH | HIGH | 19 | 10 | 55 |
| Xeros3 | Xeros/Diarizos | 0 | 0 | 485 | 97 | MODERATE | PRESCRIBED VILLAGE AREA | MODERATE | INCREASE | 22 | 7 | 15 |

**Table 3.2.**
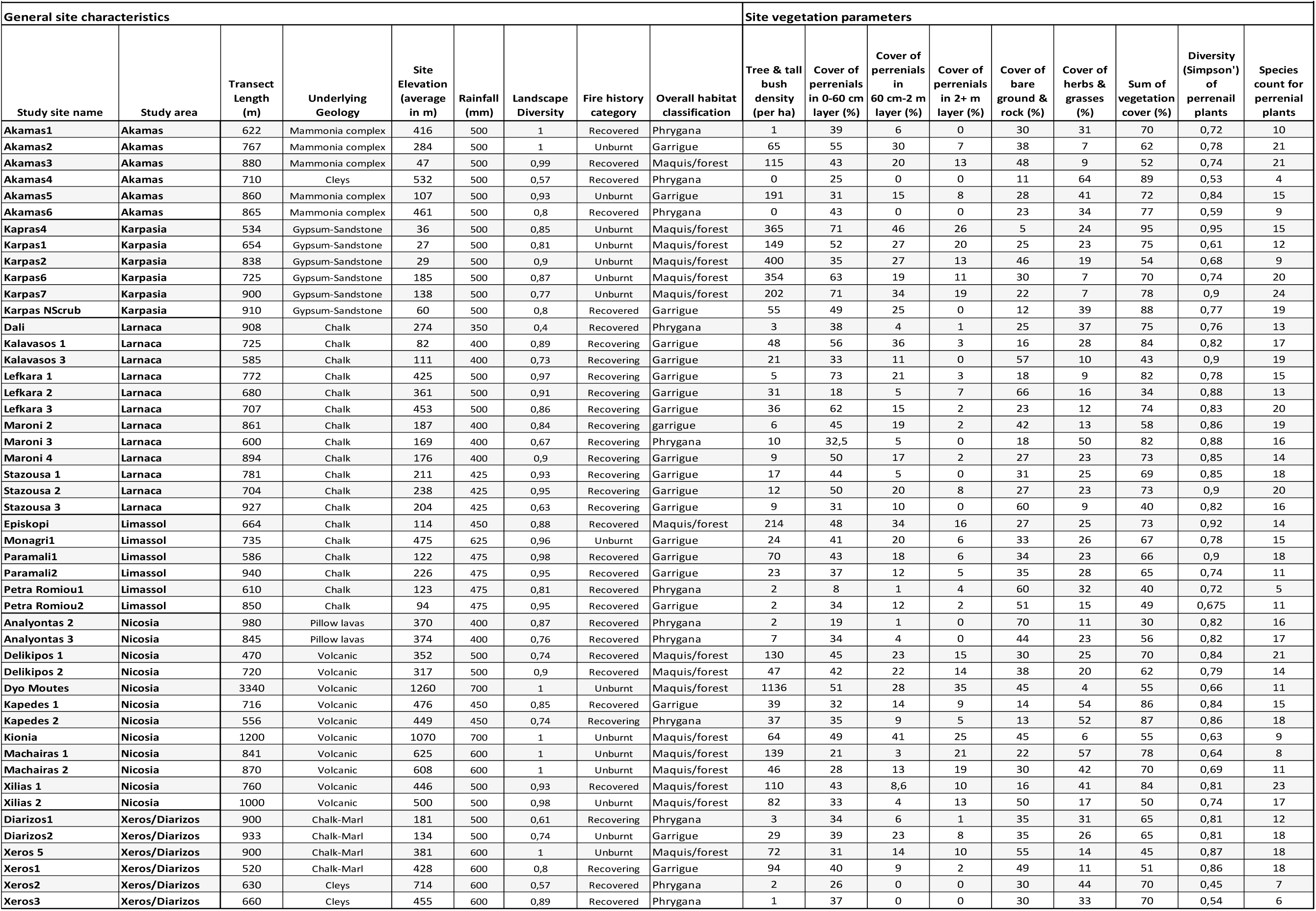
Site general characteristics and vegetation parameters (See text for explanations of how these parameters were estimated)

**Table 3.3.**
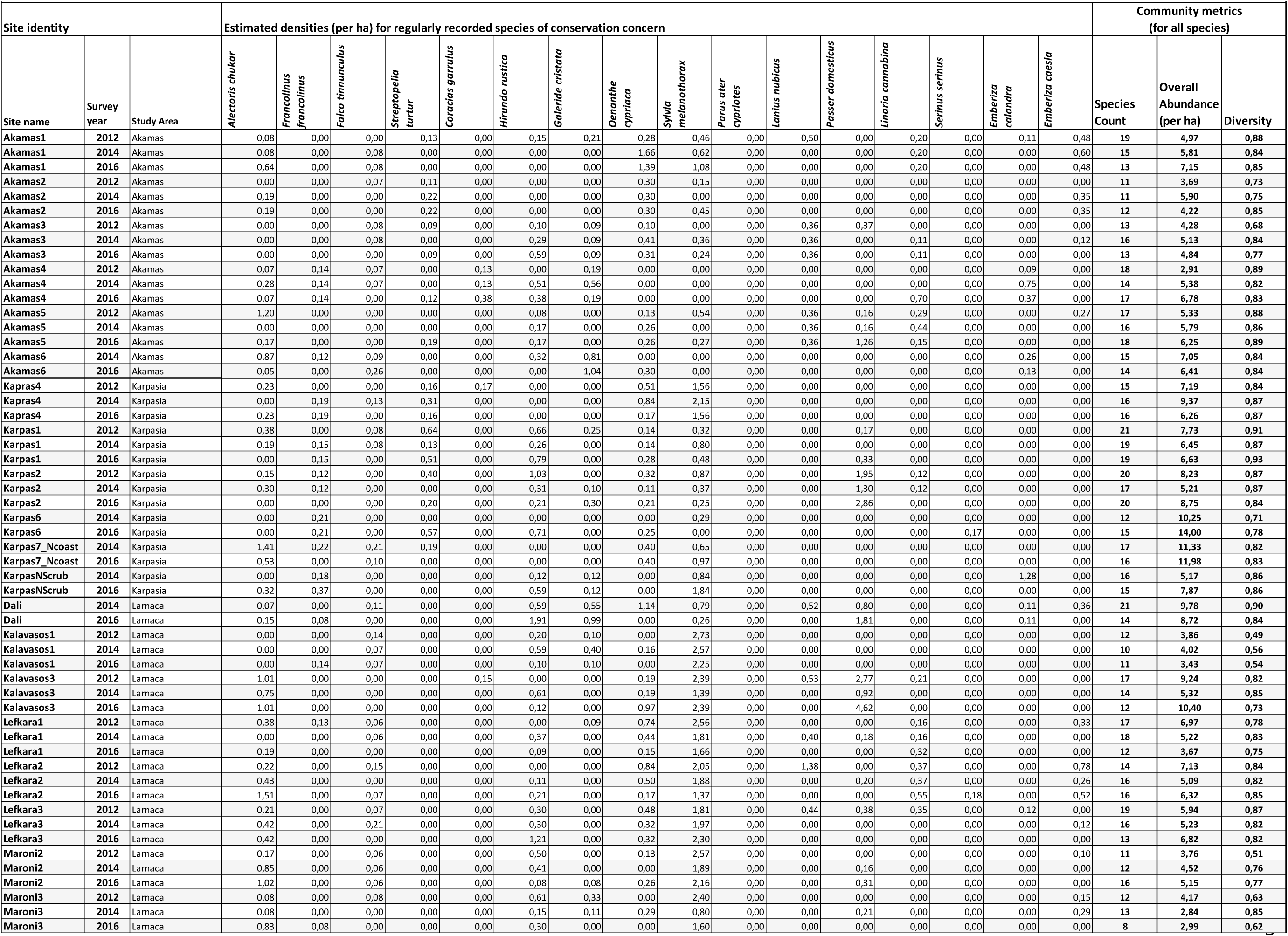
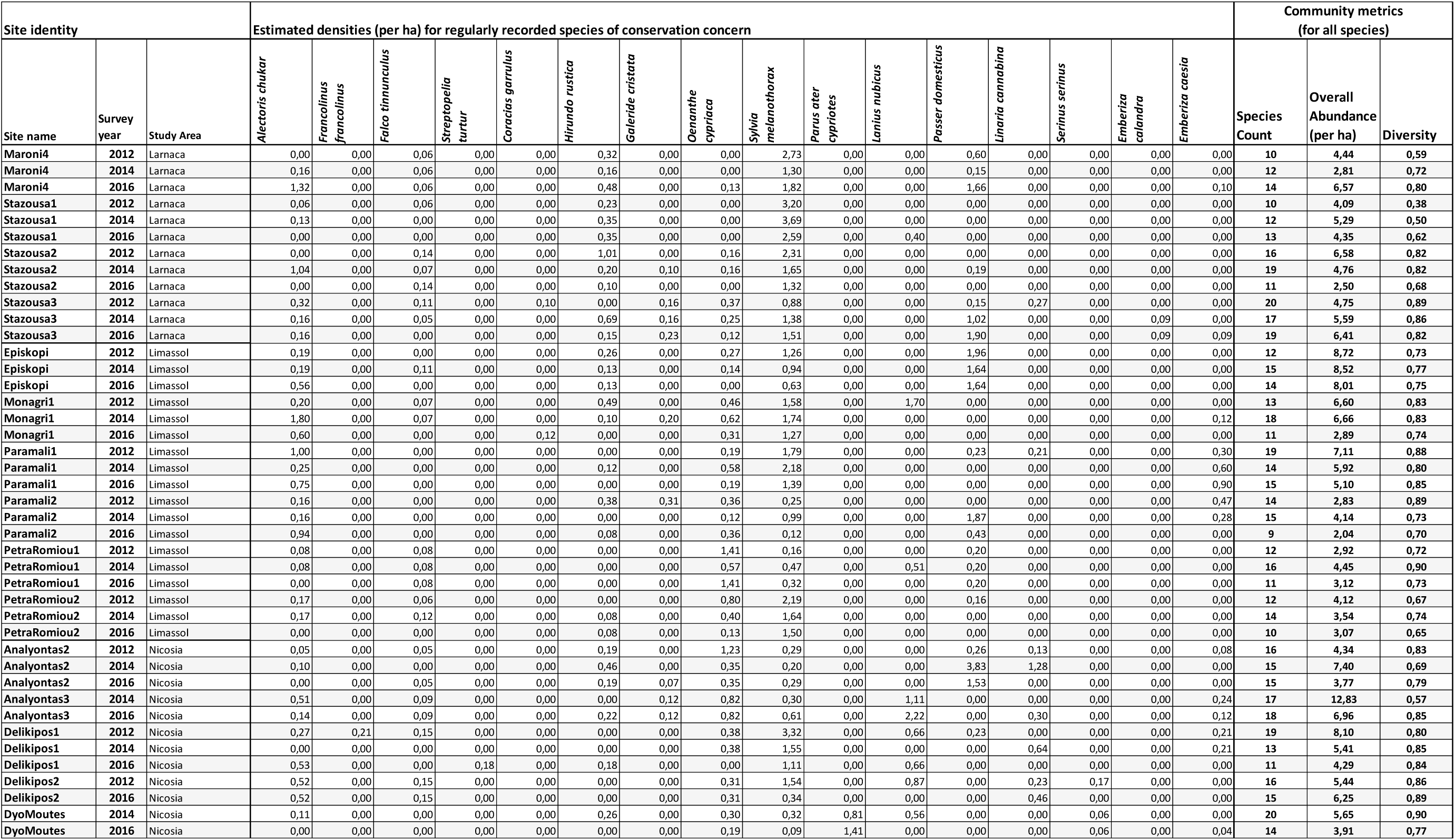
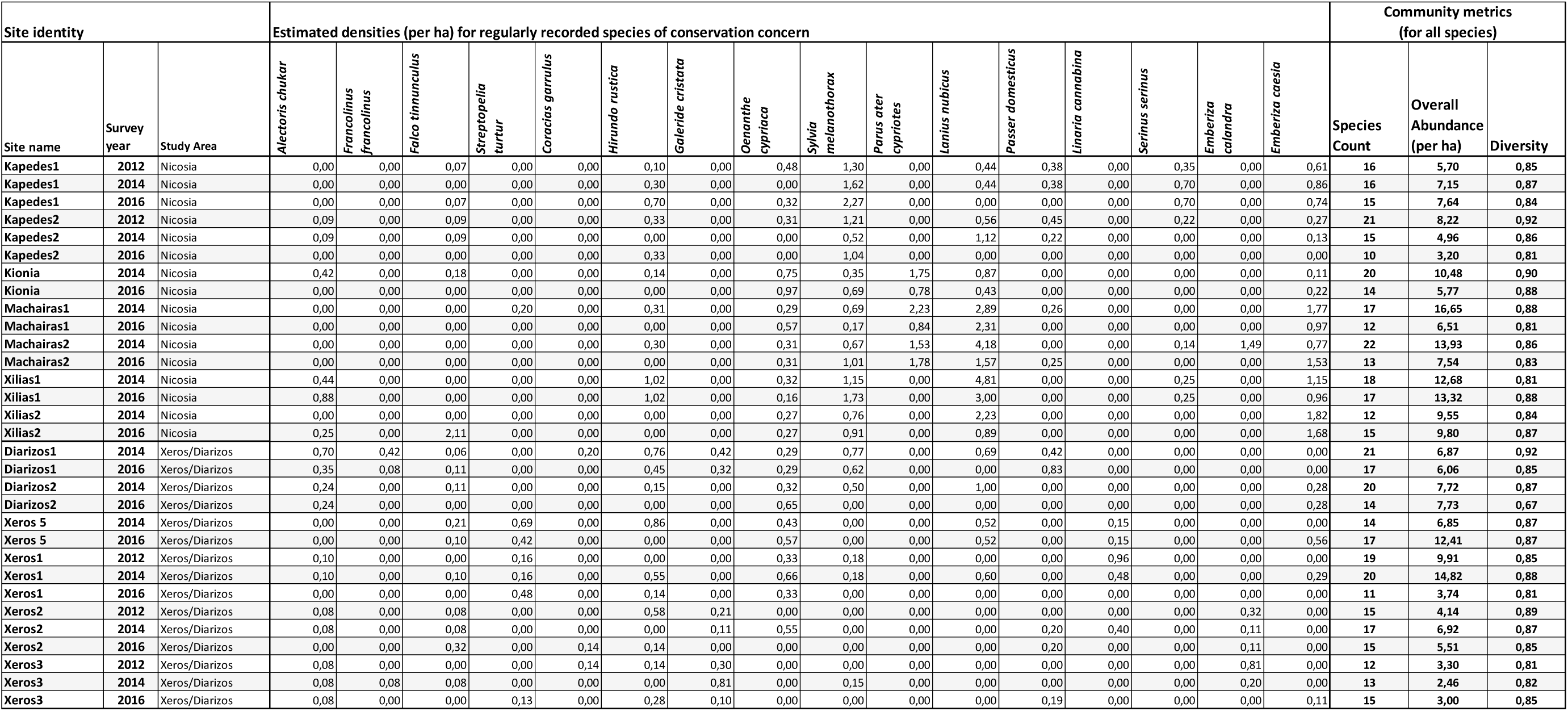
Density estimates for all regularly recorded bird species of conservation concern, plus bird community metrics for survey sites (See text for explanations of how these parameters were estimated)

## Footnotes

1 The relevant CORINE land-cover (CLC) categories were: 211 ‘Non-irrigated arable land’, 221 ‘Vineyards’, 223 ‘Olive groves’, 241 ‘Annual crops associated with permanent crops’, 242 ‘Complex cultivation patterns’, 243 ‘Land principally occupied by agriculture’.

